# *In vitro* evolution and whole genome analysis to study chemotherapy drug resistance in haploid human cells

**DOI:** 10.1101/2020.03.17.982181

**Authors:** Juan Carlos Jado, Michelle Dow, Krypton Carolino, Adam Klie, Gregory J. Fonseca, Trey Ideker, Hannah Carter, Elizabeth A. Winzeler

## Abstract

**Background:** *In vitro* evolution and whole genome analysis has proven to be a powerful method for studying the mechanism of action of small molecules in many haploid microbes but has generally not been applied to human cell lines in part because their diploid state complicates the identification of variants that confer drug resistance. To determine if haploid human cell could be used in MOA studies, we evolved resistance to five different anticancer drugs (doxorubicin, gemcitabine, etoposide, topotecan, and paclitaxel) using a near-haploid cell line (HAP1) and then analyzed the genomes of the drug resistant clones, developing a bioinformatic pipeline that involved filtering for high frequency alleles predicted to change protein sequence, or alleles which appeared in the same gene for multiple independent selections with the same compound. Applying the filter to sequences from 28 drug resistant clones identified a set of 21 genes which was strongly enriched for known resistance genes or known drug targets (*TOP1, TOP2A, DCK, WDR33, SLCO3A1)*. In addition, some lines carried structural variants that encompassed additional known resistance genes (*ABCB1, WWOX and RRM1)*. Gene expression knockdown and knockout experiments of 10 validation targets showed a high degree of specificity and accuracy in our calls and demonstrates that the same drug resistance mechanisms found in diverse clinical samples can be evolved, discovered and studied in an isogenic background.

## INTRODUCTION

In human cells, methods for discovering genes that play a role in drug resistance or which encode drug targets, especially for poorly characterized compounds, such as natural products, are limited. Genome-wide CRISPR-*Cas9* knockdown experiments[1-3] in the presence of a drug are useful to broadly implicate relevant genes, but cannot readily reveal critical gain-of-function, single nucleotide alleles, such as imatinib-resistance conferring mutations in BCR-Abl. Discovering common alleles in whole genome sequences of tumors from cohorts of patients that have relapsed after drug treatment requires very large datasets and is complicated by patient heterogeneity. Furthermore, such studies also cannot be used on experimental therapies.

Work in other organisms has shown that *in vitro* evolution and whole genome analysis (IVIEWGA) is a powerful method to discover both a comprehensive set of drug resistance alleles, as well as the targets of compounds with unknown mechanisms of action[4, 5]. In this method, clonal or near clonal organisms are isolated and then clones are subjected to increasing levels of a drug that inhibits growth. After selection, the organism is cloned again. The genomes of resistant clones are then compared to the sensitive parent clone using next generation sequencing (NGS) methods. In organisms such as *Saccharomyces cerevisiae*[6], *Plasmodium falciparum*[4, 5], Mycobacteria[7], Trypanosomes[6], and *Acinetobacter baumannii*[8] this method has been used to comprehensively discover resistance conferring variants. Surprisingly, the data shows that typically only a small number of *de novo* variants are detected after evolution. If multiple selections are performed on independent clones, the same resistance gene will appear repeatedly, although often appearing with different alleles, providing a high level of statistical confidence that the allele has not arisen by chance.

Many of the organisms on which IVIEWGA has been used with success have both haploid and diploid phases of their lifecycle, which means that selections can be performed in a haploid stage. Selecting for resistant clones in a haploid organism greatly simplifies analysis as a driver resistance allele will approach 100% frequency. In addition, for loss of function alleles, only one mutation is needed (versus both copies). Although metazoans are all diploid, haploid human cells lines are nevertheless available: HAP1, is a human chronic mylogenous leukemia (CML)-derived cell line that is completely haploid except for a 30 megabase fragment of chromosome 15 [9]. HAP1 has been used for genetic studies because mutated phenotypes are immediately exposed[10-15].

Using five different anticancer drugs (Doxorubicin, Gemcitabine, Etoposide, Topotecan, and Paclitaxel) as examples, we show that *in vitro* evolution in HAP1 cells can be used to identify the molecular basis of drug resistance in human-derived cells. Through our unbiased analysis of evolved clones, we detect a limited number of genes that acquire SNVs or CNVs after prolonged, sublethal exposure to our selected xenobiotics. We further demonstrate the power of the approach by using shRNAs and CRISPR-*Cas9* to downregulate or reintroduce selected alleles and demonstrate that this confers resistance or sensitivity to the drug which elicited the evolved genomic change. Our work has implications for clinical intervention strategies to prevent the emergence of drug resistance and tumor recurrence through gene mutations acquired through DNA damage from chemotherapeutics or natural variants which exist and persist from the heterogenous tumor cell environment.

## RESULTS

### Selection of compounds for resistance studies

To identify xenobiotics with the best efficacy against HAP1 cells we first measured ATP levels (CellTiterGlo) treating HAP1 cells with serial dilutions of 16 different drug for 48 hours. Five drugs showed EC_50_ values between 5 to 340 nM (**Fig. 1A-B, Table S1**). These included doxorubicin (DOX, EC_50_ = 46.05 ± 4.6 nM), also known as adriamycin, an anthracycline antibiotic that works by inhibiting topoisomerase II alpha (*TOP2A*)[16, 17]; gemcitabine (GEM, EC_50_ = 8.7 ± 0.7 nM), a synthetic pyrimidine nucleoside prodrug that is used against a variety of hematopoietic malignancies[18-20]; etoposide (ETP, EC_50_ = 338.6 ± 39.7 nM), a semisynthetic derivative of podophyllotoxin that also targets *TOP2A* and prevents re-ligation of the double-stranded DNA[21]; paclitaxel (PTX, EC_50_ = 17.5 ± 4.0 nM) also known as taxol, an effective anticancer agent that targets tubulin, perturbing the cytoskeleton and causing M phase cell-cycle arrest[22], and topotecan (TPT, EC_50_ = 5.6 ± 0.1 nM), a semisynthetic water-soluble derivative of camptothecin that inhibits topoisomerase I (*TOP1*)[23]. Our HAP1 EC_50_ values were similar to those previously reported for other CML cell lines (www.cancerrxgene.org [24, 25]) with the with the exception of etoposide, which appeared more effective in HAP1 cells (EC_50_ = 338.6 ± 39.7 nM) relative to other CML cell lines (> 1 µM in BV-173, KU812, EM-2, MEG-01, JURL-MK1, KCL-22, RPMI-8866, LAMA-84, K-562).

**Fig. 1.**
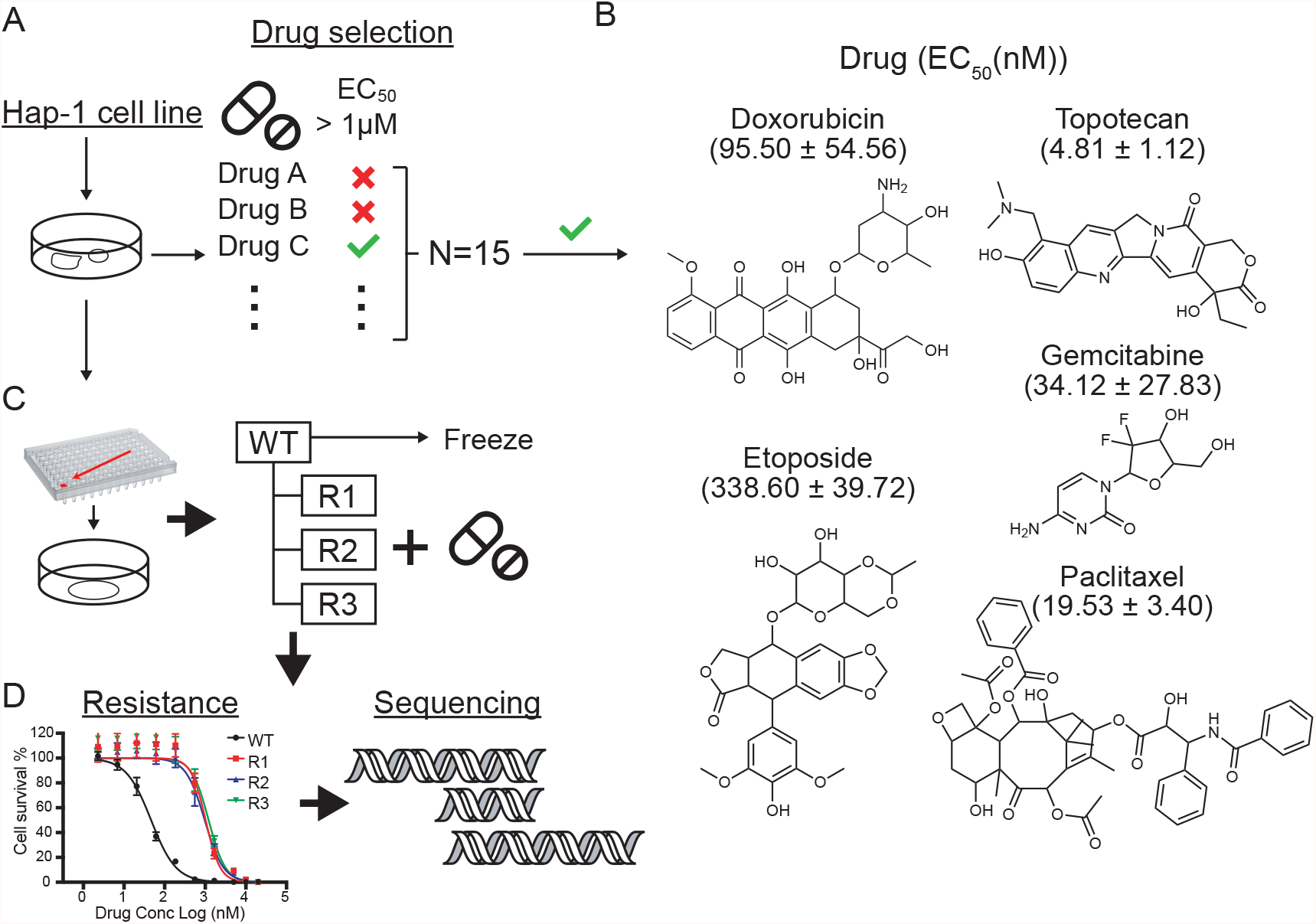
Experimental workflow. **A**. Chemotherapy drug evaluation. EC_50_ dose response assays were performed on 15 different chemotherapeutic agents (Table S1). Only drugs to which HAP1 cells were sensitive (EC_50_ value below 1µM) were considered for IVIEWGA. **B**. Chemical structures of the chemotherapy agents ultimately used for IVIEWGA. EC_50_ values are presented as the mean ± s.e.m., for n=3 biological replicates and n=8 with technical replicates per concentration point. **C**. Clone selection. To ensure a homogenous genetic background limiting dilution cloning was used to isolate individual cells prior to drug selection. For each drug three independent selections were performed. Resistance was confirmed using dose-response assays **D**. Drug resistance was achieved in 7 to 30 weeks approximately (49 and 210 generations). The parental cell line and the drug resistant lines were then sequenced.

### Evolution of resistance is readily achieved for all compounds

Our next objective was to create drug resistant lines. Although we have had difficulty creating resistant lines for some drugs in some species (“irresistibles”[26]), there is precedent for resistance to the drugs included here[27-29]. To reduce the possibility of selecting mutations that were present in a nonhomogenous population of HAP1 cells and to facilitate later genomic analysis, we first cloned the cells. This was accomplished by diluting cells to an average density of ∼0.5 cells per well in a poly-L-lysine treated 96-well plate (**Fig. 1C**) and then picking clones from wells that contained single colonies. Selections were initiated with different parent clones for the different drug-specific replicates (**Fig.1C, Fig. S1**).

To create drug resistant clones, cells were grown in tissue culture dishes (reaching 60-80% semi-confluence) in the presence of sublethal concentrations of each drug using one of two different methods. Most cell lines (DOX, GEM, TPT and PTX resistant clones) were subjected to a lethal concentration (∼3-5 × EC_50_ value), killing more than 95% of the cells. Then, treatment was removed until cells reached semi-confluence again (doubling every 22 hours[30]) whereupon drug at ∼ the EC_95_ value was reapplied. Alternatively, for ETP-resistant clones a step-wise selection method was used whereby cells were repeatedly exposed to a concentration that killed around 50% of the cell population. Drug concentration was increased by 5-10% every 5 days while keeping the growth rate at 50% of untreated culture. Although others have used mutagenesis [31], we have found that this can increase the rate of background mutations, which would complicate an already difficult analysis. Because mutations will arise randomly during long term cell culture, we attempted at least three independent selections for each drug, in each case starting with an identical parental isolate (**Fig. 1C**). In a few cases, independent selections could not be achieved and dependent clones with a shared lineage (DOX-R4a and DOX-R4b; PTX-R2a and PTX-R2b; TPT-R4a, TPT-R4b and TPT-R4c) were collected. Resistance emerged after several months depending on the drug and the method used (7-30 weeks approximately, 49-210 generations) (**Fig. S1**).

Once resistance was observed in the batch culture, we isolated clones from batch drug-selected cultures and the drug sensitivity of the clone was measured and compared to the isogenic parent clones (**Fig. 1D**). We observed an EC_50_ fold shift between 3.3 and 411.7 (**Table S2**) in paired comparisons. To demonstrate that the drug resistance phenotype was stable, drug pressure was removed for 8 weeks (approximately 56 generations) and clones were retested for sensitivity. We observed no changes in the EC_50_ values, indicating that resistance phenotypes were not due to transient adaptation.

### Identification of putative resistance variants using next-generation sequencing

We next performed whole genome and exome paired-end read sequencing on the 35 cell lines (both drug-resistant clones and their matched drug-sensitive parent clones). Our IVIEWGA studies in *Plasmodium*[5], have shown that stable drug resistance is typically conferred by SNVs in coding regions and thus exome sequencing was an efficient mechanism to find causal variants. However, gene amplifications, which contribute to 1/3 of drug resistance events in *Plasmodium*[5], are more accurately detected with WGS because exact chromosomal recombination sites, which may fall in intergenic regions, can be reconstructed from WGS data. Because of falling costs over the course of the project, more samples (N=21) were ultimately whole genome sequenced than whole exome sequenced (N=14).

Sequencing quality was high for all samples: alignment showed that, on average, 99.9% of 700 million WGS (40 million WES) reads mapped to the hg19 reference genome with 86% of the bases covered by 20 or more reads (**Table S3**). By comparing sequences of evolved clones to their respective parental clones, we discovered a total of 41,259 SNVs (**Table S4**), of which 26,625 were unique (**Table S5**, Methods). The majority of variants in all cell lines was non-coding (**Table S4, S5)** and were evenly distributed with respect to chromosome length (**Fig. S2**). Of the 26,625 mutations almost all (26,468) were present at allele frequencies (AF) of less than 85% relative to their parent clone and would thus not be expected to be driver mutations, given that the parents were cloned (to the best of our ability) before selections were initiated. The five drugs varied in the number of mutations, with TPT having the highest overall numbers (**Table 1**).

**Table 1.** Summary of average number of mutations. Number of selections performed with the drug is given by n. SNVs and Indels were grouped according to snpEff sequence ontology annotations (Methods, Table S4), and detailed counts per clone can be found in Table S5.

|  |  |  | **WGS** |  |  |  | **WES** |  |  |
| --- | --- | --- | --- | --- | --- | --- | --- | --- | --- |
|  | DOX<br>(n=3) | GEM<br>(n=3) | PTX<br>(n=4) | TPT<br>(n=3) | TPT<br>(n=3) | DOX<br>(n=3) | ETP<br>(n=3) | GEM<br>(n=3) | PTX<br>(n=3) |
| **Indels** |  |  |  |  |  |  |  |  |  |
| *Disruptive inframe ins.* | 0.00 | 0.00 | 0.00 | 0.00 | 1.00 | 1.00 | 0.00 | 0.00 | 0.00 |
| *Frameshift* | 0.00 | 1.00 | 1.33 | 2.33 | 0.00 | 1.00 | 1.00 | 1.00 | 1.00 |
| *Frameshift plus stop-gained* | 0.00 | 0.00 | 0.00 | 0.33 | 0.00 | 0.00 | 1.00 | 0.00 | 0.00 |
| *Inframe insertion* | 0.00 | 0.00 | 1.00 | 0.33 | 0.00 | 0.00 | 1.00 | 0.00 | 0.00 |
| *Intergenic* | 27.67 | 43.00 | 26.75 | 24.67 | 47.67 | 2.00 | 3.50 | 1.00 | 1.00 |
| *Intragenic* | 10.00 | 5.67 | 9.00 | 9.00 | 14.33 | 1.00 | 2.00 | 1.00 | 1.00 |
| *Intron* | 12.00 | 20.33 | 16.25 | 15.67 | 32.33 | 1.00 | 1.00 | 4.00 | 1.50 |
| *Splice region plus intron* | 0.00 | 0.00 | 0.00 | 103.3 | 0.00 | 0.00 | 0.00 | 1.00 | 0.00 |
| **SNVs** |  |  |  |  |  |  |  |  |  |
| *Disruptive inframe del.* | 0.00 | 1.00 | 1.00 | 0.67 | 0.00 | 1.00 | 0.00 | 1.00 | 1.33 |
| *Frameshift* | 1.00 | 2.00 | 1.00 | 22.33 | 2.00 | 3.00 | 1.67 | 1.00 | 3.00 |
| *Inframe deletion* | 0.00 | 0.00 | 1.00 | 0.00 | 0.00 | 0.00 | 2.00 | 0.00 | 0.00 |
| *Intergenic* | 898.33 | 1303.3 | 1416.6 | 258.67 | 2635.7 | 25.00 | 19.00 | 17.00 | 21.33 |
| *Intragenic* | 272.00 | 403.33 | 389.25 | 77.67 | 834.67 | 15.33 | 12.33 | 10.33 | 11.33 |
| *Intron* | 448.67 | 701.33 | 764.00 | 128.3 | 1358.33 | 28.00 | 27.00 | 33.00 | 22.67 |
| *Missense* | 16.00 | 14.00 | 12.75 | 7.00 | 34.33 | 15.00 | 19.67 | 21.67 | 15.67 |
| *Others* | 1.00 | 1.00 | 0.00 | 0.00 | 1.33 | 1.00 | 1.00 | 1.50 | 1.00 |
| *Splice region plus intron* | 1.00 | 1.67 | 2.00 | 30.00 | 3.33 | 1.67 | 1.50 | 1.00 | 1.33 |
| *Start lost* | 0.00 | 0.00 | 0.00 | 0.00 | 0.00 | 0.00 | 0.00 | 1.00 | 1.00 |
| *Stop gained* | 0.00 | 2.50 | 1.00 | 0.00 | 1.67 | 1.50 | 2.00 | 2.67 | 0.00 |
| *Stop lost* | 0.00 | 1.00 | 0.00 | 0.00 | 0.00 | 0.00 | 0.00 | 0.00 | 0.00 |
| *Synonymous* | 3.00 | 7.33 | 4.00 | 6.33 | 7.67 | 6.67 | 5.33 | 7.67 | 5.33 |

We next developed a pipeline (**Fig. S3A**, Methods) to filter the 26,625 “called” mutations (**Table S6**) to a final list of potential variants conferring drug resistance (**Table 1**). Our previous analyses in other species suggested that variants presented in coding regions are more likely to contribute to drug resistance even though this could exclude the variants associated with certain transcription factor (TF) binding sites. Therefore, our strategy focused on mutations that were in exonic regions and were drug-specific (**Fig. S3A**). We further considered only mutations likely to have a functional impact at the protein level (missense, nonsense, frameshift, start or stop gain or loss) which further reduced the number to an average of 35 and 23 nonsynonymous mutations for WGS and WES, respectively (**Fig. S3A**). Reasoning that resistance driver mutations (e.g. those actually causing resistance) would be present in 100% of the haploid cells in the sequenced culture, we selected only the variants with high allele frequency (AF > 0.85, as determined by sequencing read count). The top 2.5% of highest AF mutations corresponded to an AF > 0.85 (**Fig S3B**). At this cutoff, the majority of cell lines harbored a candidate resistance mutation. While selecting a cutoff represents a tradeoff with potentially missed relevant mutations, the full list of mutations is provided in the supplement (**Table S6**). We did not note any strong correlation between read depth and allelic fraction in our study (R2 = 0.06; **Fig. S3C**) and all of the final mutations selected for further analysis had a read depth > 10 reads, with the majority supported by over 20 (**Fig. S3D**). Although some with AF <0.85 could confer a beneficial advantage to the cell, most are likely to be random mutations that arose during long term culture. Finally, based on our experience with microbes whereupon genes with multiple, predicted independent amino acid changes (not expected by chance (4 genes, STARD9, CYP1B1, SLCO3A1 and DCK)) are often found for the same drug, we added these genes to our final list of 21 candidates (**Table S7**).

### Somatic Copy number variations (CNVs)

We next searched for CNVs (both amplifications and deletions) in our WGS and WES data using Control-FreeC[32]. Overall patterns of broad and focal alterations across the drugs and conditions varied (**Fig. S4A, Table S8**). Using a corrected p-value of less than 0.05, we identified 93 total amplification and 108 deletion events, with most appearing in the TPT-resistant samples (123) (**Table S8**). The CNVs had an average size of 8.5 Mbp (stdev 19 Mbp), ranged from 15,000 bp to 152 Mbp (**Fig. S4A**) and covered ∼3% of the genome, on average. More CNVs were called in WES samples because of sequencing gaps—even for WGS samples, some CNVs were separated by short distances and were nearly contiguous (**Fig. S4A**). It is likely that some CNVs were also missed in the WES data. The number of events was proportional to chromosome size, with the exception of the Y chromosome, for which there were ∼4 × more events (47) per unit length. Some CNV calls were supported by paired end red data, for example, the one near WWOX (**Fig. S4B, C**).

### Doxorubicin resistance is associated with mutations in *TOP2A* and a solute carrier transporter

To evaluate the approach, we next considered the set of SNVs and CNVs for each drug. For DOX, we had six available selections from two different starting clones (WT-1 (WGS) and WT-5 (WES)) that were analyzed by WGS (DOX-R1, DOX-R2, DOX-R3) and by WES (DOX-R4a, DOX-R4b and DOX-R5) (**Fig. 2A**). High allele frequency missense mutations were found in only 11 genes (**Table S7**). Of note, DOX-R2 and DOX-R3 carried mutations in *TOP2A* at frequencies of 0.89 and 0.87, respectively. This is the known target of DOX[21, 33] and is known to play a role in drug resistance[33-35]. The amino acid mutation, Pro803Thr (**Fig. S5**), sits within the principal DNA-binding locus, the DNA-gate, a region conserved in type II topoisomerases (*TOP2A* and *TOP2B*). It is also adjacent to the catalytic tyrosine (Tyr805), responsible for nucleophilic attack on DNA[36]. While one explanation is that Pro803Thr creates steric hindrance and blocks DOX access to the site, a more likely explanation is that the mutation is a loss-of-function mutation, especially as knockdown of TOP2A activity has previously been shown to confer DOX resistance in a Eμ-Myc mouse lymphoma system[37]. To reproduce these results in our HAP1 human cells, *TOP2A* was downregulated using a shRNA pool containing constructs encoding target-specific shRNA hairpins for *TOP2A*. Western blots further showed the expected down regulation of protein levels (**Fig. 2B**) and an EC_50_ analysis of the wildtype and the knockdown line revealed a 4.25-fold increase in DOX resistance compared to the isogenic parent (**Fig. 2C, D**).

**Fig. 2.**
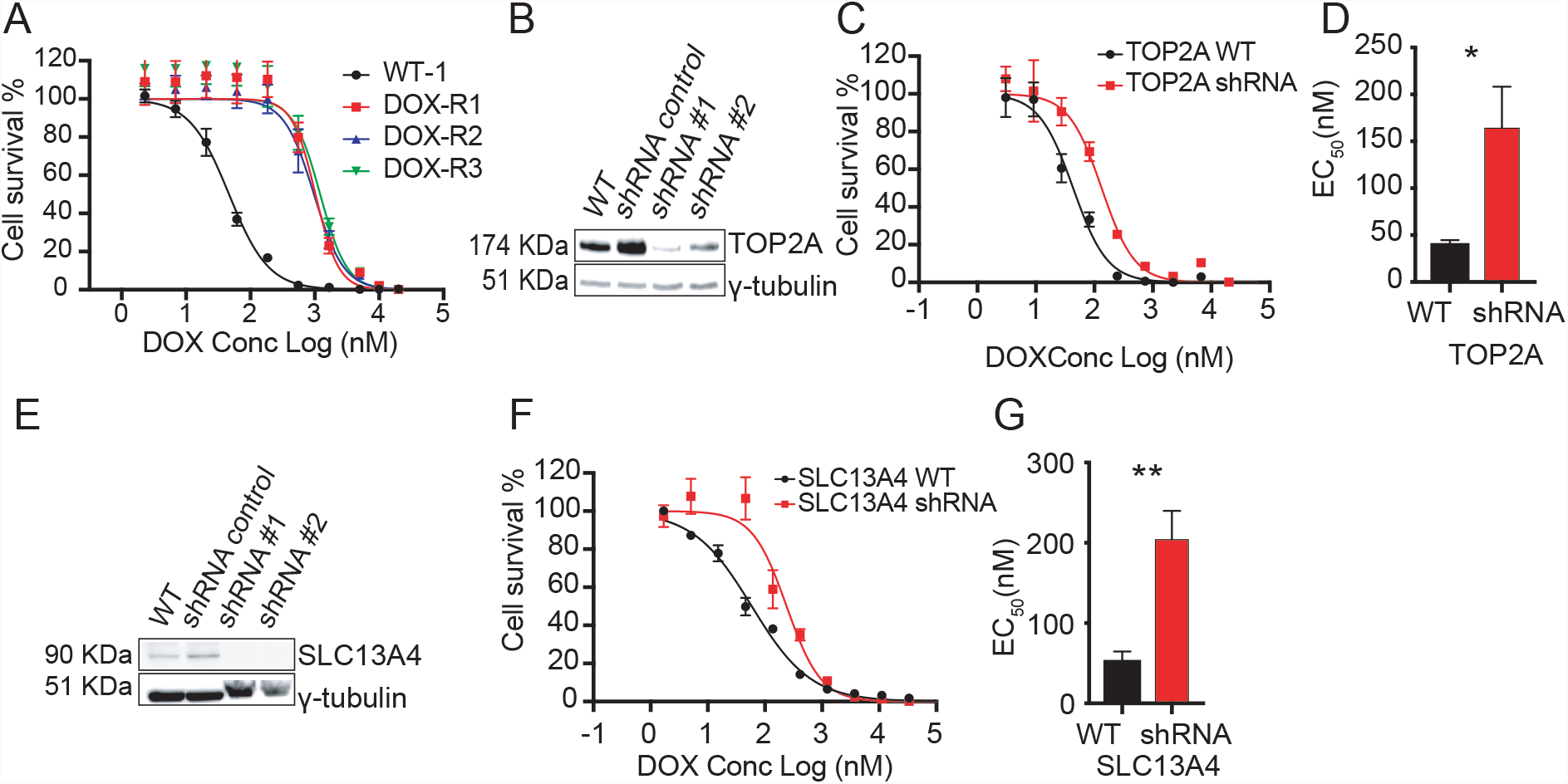
**A**. DOX EC_50_ curves for DOX evolved clones using 8 technical replicates for each concentration **B**. Western blot confirming that shRNA gene depletion downregulates TOP2A protein level. **C**. EC_50_ curves of the WT and shRNA (shRNA #1) knockdown cell lines for *TOP2A*. **D**. Barplot of the WT and shRNA knockdown cell line (shRNA#1) for *TOP2A*. **E**. Western blot confirming protein downregulation of SLC13A4 by shRNA. **F**. EC_50_ curves of the WT and shRNA knockdown cell lines for *SLC13A4*. **G**. Barplot of the WT and shRNA knockdown cell lines for *SLC13A4*. Data is represented by mean ± s.e.m. with n=2 biological replicates and n=8 technical replicates (*TOP2A*) or n=3 biological replicates and n=4 technical replicates (*SLC13A4*) for every concentration point. * = p value < 0.05, ** = p < 0.01. p values determined by two-tailed *t* test.

We also found missense mutations present in 100% of the reads for several other attractive but less well characterized genes; *SLC13A4* (Gln165His, DOX-R4b), and *SPG7* (Lys593Asn, DOX-R5), as well as one uncharacterized gene (AC091801.1, His13Asn, DOX-R4a) in the three different clones that were subjected to WES and were derived from WT-5. *SLC13A4* is a solute carrier transport family member and members of this general solute carrier family have appeared in selections conducted in microbes (e.g. the UDP-galactose transporter and the AcetylCoA transporter[38]) and are also associated with cancer drug resistance[39]. The Gln165His mutation is located in the disordered region of the protein. To validate *SLC13A4* we performed a gene knockdown using a shRNA pool that targeted *SLC13A4*, containing three expression constructs each encoding target-specific 19-25 nucleotide shRNA hairpins. Protein expression levels of the knockdown line were measured by western blot followed by a dose-response assay to compare its EC_50_ value with the wildtype line (**Fig. 2E**). The 4 × increase in resistance suggests that *SLC13A4* contributes to resistance, although it may not account completely for the level of resistance of the sequenced clones, which ranged from 4 to 11 × (**Fig. 2F, G**).

### Gemcitabine resistance is conferred by changes in DCK and RRM1 activity

Six selections were performed with GEM (starting from two different isogenic parents; WT-2 (WGS) and WT-3 (WES)). Among those, three GEM-resistant clones subjected to WGS (GEM-R1, GEM-R2 and GEM-R3) showed an average EC_50_ shift of 300 to 400-fold (**Fig. 3A, Table S2**), and the clones showed no change in HAP1 sensitivity to other drugs (**Fig. 3B**). As there were no candidate alleles with AF > 0.85, we looked for genes that acquired mutations in multiple selections, identifying deoxycytidine kinase (*DCK*) as likely important for resistance. Interestingly, across cell lines several distinct mutations were found in *DCK*, with varying effects (missense and frameshift) across several different positions (**Table 2**). In particular, the missense substitution Ser129Tyr, present in GEM-R1 and GEM-R3, not only alters the amino-acid size and charge also falls at the end of exon 3, within the ATP-binding pocket of a phosphorylation site, making it a strong causal candidate for GEM drug resistance (**Fig. S6**). GEM only becomes pharmacologically active if it is phosphorylated and the first phosphorylation is catalyzed by DCK. A shRNA knockdown of DCK was performed and confirmed by western blot analysis (**Fig. 3C**). Downregulation of the gene resulted in a 36.5-fold increase in the EC_50_ value compared to both the isogenic parent line and the shRNA negative control (**Fig. 3D, E; Table 2**).

**Table 2.** Validation (knockdown) results for selected genes. CNV, copy-number variant. MS, missense. FS, frameshift variant, KO/KD/CI, knockout, knockdown, chemical inhibition (verapamil, ABCB1). ND: No data: gene knockdowns were attempted but could not be achieved. NE: Not expressed (protein not detected by Western blot, preventing validation). *EC*_*50*_ *WT and EC*_*50*_ *KO/KD/CI* are from matched pairs for the given drug and represent the mean ± s.e.m. (n=3 biological replicates).

| *Drug* | *Sample* | *Gene* | *Amino Acid Change* | *Type* | *AF* | *EC<sub>50</sub> (nM) WT* | *EC<sub>50</sub> (nM) KD/KO/CI* |
| --- | --- | --- | --- | --- | --- | --- | --- |
| **DOX** | R1, R2, R3 | TOP2A | Pro803Thr | MS | 0.89, 0.87 | 38.6 ± 4.3 | 164.3 ± 43.9 |
|  | R4b | SLC13A4 | Gly165His | MS | 1 | 52.9 ± 11.6 | 204.3 ± 35.7 |
|  | R5 | SPG7 | Lys593Asn | MS | 1 | NE* |  |
| **PTX** | R1, R2a, R3 | WWOX | 16q23.1 | CNV | - | 5.8 ± 2.5 | 42.9 ± 11.7 |
|  | R2a, R2b | ABCB1 | 7q21.12 | CNV | - | 252 ± 38<br>218 ± 14 | 1.3 ± 0.1<br>1.3 ± 0.1 |
|  | R2b, R6 | SLCO3A1 | Ile587Asn (R2b),<br>Ala263Thr (R6) | MS |  | 6.6 ± 3.3 | 51.5 ± 9.9 |
| **GEM** | R1, R2, R3 | DCK | Ser129Tyr (R1, R2),<br>Asn80fs (R1, R3);<br>Asn113fs (R2);<br>Thr184fs (R3) | MS,<br>FS |  | 14.3 ± 1.7 | 521.7 ± 58.3 |
|  | R4, R5, R6 | RRM1 | 11p15.4 | CNV | - | 54.9 ± 5.8 | 1.8 ± 0.1 |
| **ETP** | R2, R3 | WNT3A | 1q42.13 (R2) | CNV | - | ND | - |
|  | R3 | WDR33 | P622T | MS | 1 | 241.5 ± 31.0 | 821.6 ± 226.9 |
| **TPT** | R1, R4a, R4b, R4c | CYP1B1 | Val432Leu;<br>Asp217Glu (R4a,b,c) | MS | 0.13, 0.40,<br>0.43, 0.42 | 6.3 ± 0.2 | 13.3 ± 0.3 |
|  | R1, R2, R3 | WWOX | 16q23.1 | CNV | - | 2.4 ± 0.3 | 22.8 ± 2.7 |
|  | R4a, R4b, R4c | TOP1 | His81fs; 20q12 | FS;<br>CNV | 1 | ND | - |
|  | R4a, R4b, R4c | USP47 | Arg408* | Stop | 0.38, 0.57,<br>0.58 | 3.05 ± 0.2 | 1.07 ± 0.07 |

**Fig. 3.**
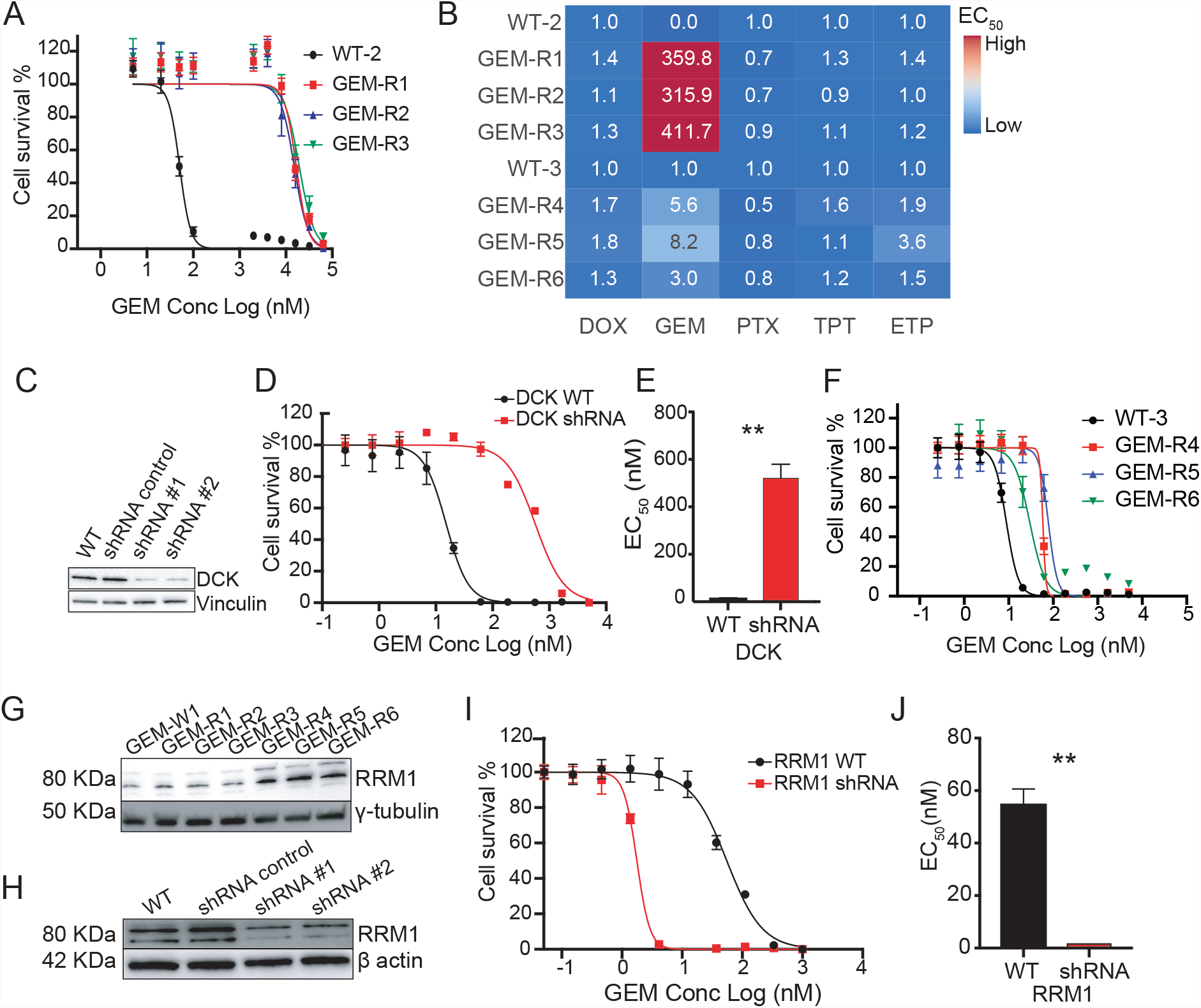
**A**. GEM EC_50_ for curves first set of GEM evolved lines using n=8 technical replicates per concentration point **B**. EC_50_ ratio matrix showing absence of multidrug resistance pathways of GEM resistant lines. **C**. Western blot confirming that shRNA gene depletion downregulates protein levels for *DCK*. **D**. EC_50_ curves of the WT and shRNA knockdown cell lines for *DCK*. n=8 with individual technical replicates overlaid for every concentration point. **E**. Barplot of the WT and shRNA knockdown cell lines for *DCK*. **F**. GEM EC_50_ curves for second set of GEM evolved lines. **G**. Western blot for *RRM1* across all GEM samples showing overexpression pattern of *RRM1* in GEM-R4-6 resistant clones. *γ-tubulin* is used as a loading control. **H**. Western blot confirming that shRNA gene depletion downregulates protein levels for *RRM1. β-actin* is used as a loading control. **I**. EC_50_ curves of the WT and shRNA knockdown cell lines for *RRM1*. n=4 with individual technical replicates overlaid for every concentration point. **J**. Barplot of the WT and shRNA knockdown cell lines for *RRM1*. All data is represented by mean ± s.e.m. with n=3 with individual biological replicates overlaid. ** = p value < 0.01. p values determined by two-tailed *t* test.

The three WT-3 derived GEM-resistant clones (GEM-R4, GEM-R5 and GEM-R6) subjected to WES were not as resistant as those used in WGS (∼6 × versus ∼400 ×, **Fig. 3F, Table S2**). Our work in other species with well characterized compounds suggests this is not surprising and that even single nucleotide changes in the same gene can yield different levels of resistance. For example, repeated selections with dihydroorotate dehydrogenase (DHOD) inhibitors in a mouse model and *in vitro* culture gave rise to 13 different point mutations in parasite DHODH with levels of resistance ranging from 2-to ∼400-fold [40]. No high AF SNVs were evident in these lines and *DCK* exons were not mutated. On the other hand, the three WES clones contained 20 CNVs that could play a role in drug resistance. Most CNVs were not shared between lines but GEM-R4, GEM-R5 and GEM-R6 all bore overlapping CNVs of varying sizes on chromosome 11, with all three lines bearing 3-4 copies (p value = 1.38e-37 to 2.05e-142) (**Fig. S4**). The chromosome 11 CNV was only found in GEM resistant lines and not in any of the other evolved lines (in contrast to CNVs on chromosome 1 or 16, for example). While it is difficult to determine which of the 140 genes in the smallest interval contribute to resistance, a known resistance mediator or target of GEM, ribonucleotide reductase (*RRM1*), was found within the amplified region. RRM1 is the largest catalytic subunit of ribonucleotide reductase, a key enzyme catalyzing the transformation of ribonucleotide diphosphates to deoxyribonucleoside diphosphates that are required for DNA synthesis and repair, and GEM is known to inhibit DNA polymerase by inhibiting *RRM1*[41]. Furthermore, overexpression of RRM1 is associated with poorer prognosis after gemcitabine treatment in non-small cell lung cancer[42] and in bladder cancer[43].

Western blot analysis of the evolved lines showed that the amplification was indeed associated with increased protein levels (**Fig. 3G**). As an additional validation, we performed a single shRNA knockdown of *RRM1* to reduce protein expression (**Fig. 3H**), followed by a dose-response assay comparing EC_50_ values of both wildtype HAP1 and *RRM1* knockdown lines, which showed that downregulation of *RRM1* made HAP1 cells 31-fold more sensitive to GEM than their isogenic parent (**Fig. 3I, J**). As expected *RRM1* downregulation had no effect on HAP1 sensitivity to other drugs (**Fig. S7**).

### Etoposide resistance is modulated by levels of *WDR33*

We created three independent ETP resistant clones, all of which were subjected to WES, and compared them to one isogenic parent clone (WT-3) (23, 13 and 9-fold increased resistance respectively (**Fig. S8A, Table S2**). A single gene, *WDR33* (ETP-R3), carried a SNV (Pro622Thr) with a 100% allele frequency. This gene encodes for a member of the *WD* repeat protein family and is one of the six subunits of a multiprotein complex called *CPSF* (cleavage and polyadenylation specific factor)[44] involved in cell cycle progression, signal transduction, apoptosis and gene regulation. Disruption of *WDR33* can lead to slowed DNA replication forks[45], which could potentially explain why its disruption protects against topoisomerase inhibitors that block DNA unwinding. Lines in which *WDR33* was knocked down via shRNA acquired an EC_50_ value 3.4 times greater than its parental line or the scrambled control (**Fig. S8B-D; Table 2**), despite an incomplete disruption of the gene by shRNA silencing.

No clear candidate SNVs were evident for ETP-R1 and ETP-R2, which did not carry the *WDR33* mutation (**Table S6, Table S7**). All ETP lines carried multiple CNVs, however, including a large shared amplification on chromosome 15 (ETP-R1 and R3). Approximately 120 protein coding genes are found in this region, including *BUB1B*, the BUB1 mitotic checkpoint serine/threonine kinase B, *BMF*, a BCL-modifying factor, as well as the *RAD51* recombinase, whose overexpression has been previously shown to confer ETP resistance[46]. Overexpression of *RAD51* activity sensitizes cells to a variety of cancer drugs, including etoposide, doxorubicin and topotecan[47]. Notably, all ETP resistant lines were also cross-resistant to PTX, TPT and DOX, providing support for this general mechanism of resistance.

### Paclitaxel resistance is mediated by transporters *SLCO3A1 and ABCB1*

Seven different paclitaxel lines were created with different resistance levels (PTX-R1, R2a, R2b and R3, ∼10 × to PTX-R4, R5, R6, 50X) (**Table S2**). The first four (**Fig. 4A**) were subjected to WGS and the latter three to WES. SNV analysis yielded no candidate genes (frameshift, indels, and missense mutations with an allele frequency >0.85). From genes with an allele frequency of less than 0.85, *SLCO3A1*, encoding another solute carrier transporter, was notable in that multiple missense alleles were identified (Ile587Asn, Ala263Thr). This class of transporter is known to play a role in the import of drugs as well as hormones such as prostaglandin[48]. Gene knockdown experiments showed that clones with loss of SLCO3A1 (**Fig. 4B**) resulted in HAP1 cells that were ∼8 times more resistant than their isogenic parents to PTX (**Fig. 4C, D**).

**Fig. 4.**
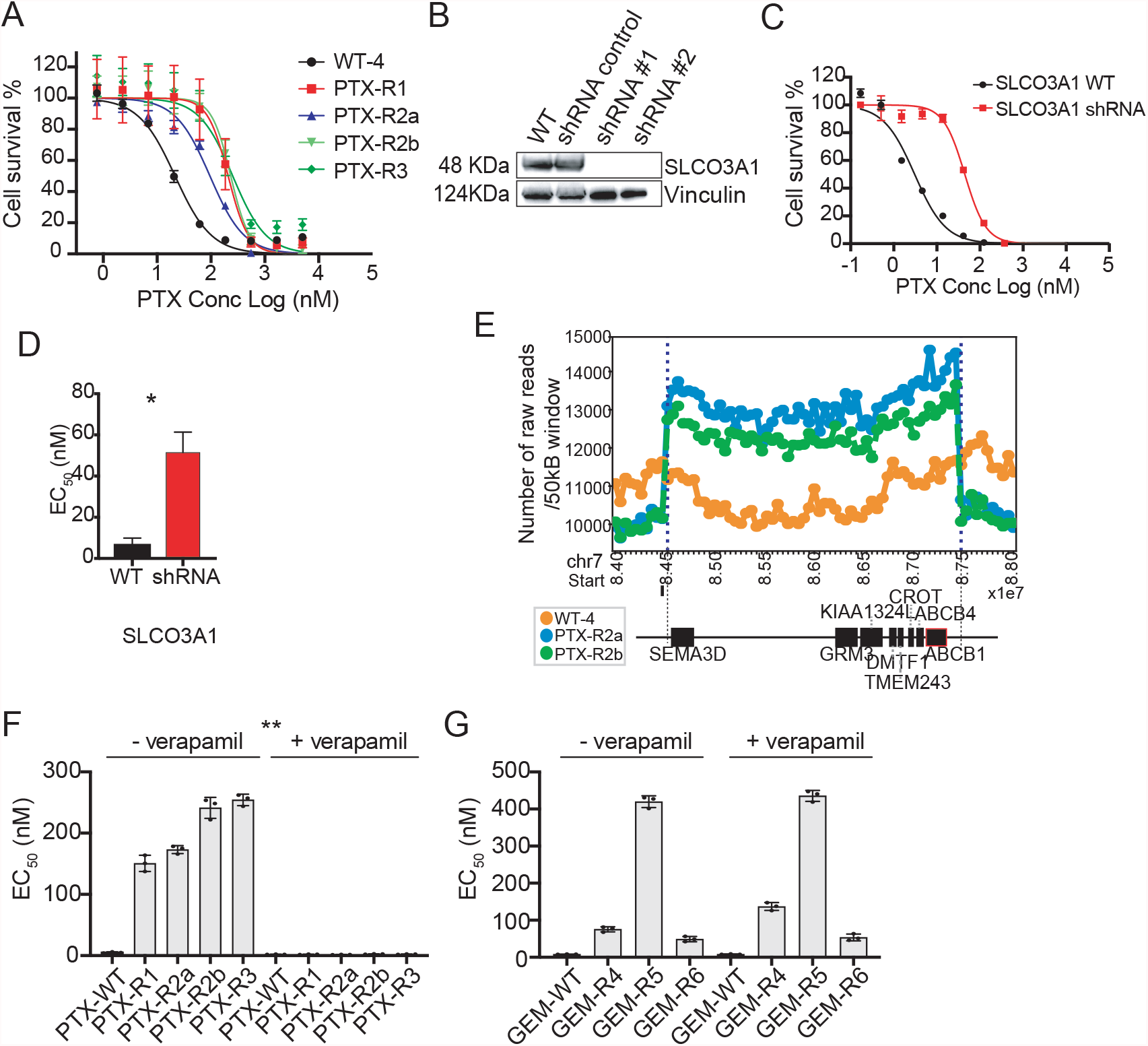
**A**. PTX EC_50_ curves for evolved lines with n=8 technical replicates (values are in Table S2) **B**. Western blot confirming shRNAs downregulate protein levels for *SLCO3A1*. **C**. EC_50_ curves of the WT and shRNA knock-down cell lines. **D**. Barplot of the WT and shRNA knockdown cell lines for *SLCO3A1*. **E**. Raw copy number profile for the amplification event containing protein coding genes including ABC transporters (*ABCB1/ABCB4)* PTX cell lines. The amplification region (chr7:84,500,000-87,300,000) had a higher number of raw reads (labeled with blue dash lines) with default window size of 50K bp. Genes associated with the CNV event are depicted by black boxes underneath according to their position and sizes. *ABCB1* is highlighted with red outline. **F**. Barplot of EC_50_ of the PTX treated cell lines with and without verapamil and verapamil alone showing sensitization in presence of verapamil as ABC inhibitor (n=4 technical replicates). **G**. Barplot of EC_50_ of the GEM cell lines ± verapamil showing no EC_50_ shift for GEM cell lines when co-treated with verapamil. Unless otherwise noted, all data is represented by mean ± s.e.m. with n=3 with individual biological replicates overlaid. ** = p value < 0.01. p values determined by two-tailed *t* test.

Despite the lack of obvious coding SNVs, PTX-R1, R2a, R2b and R3 had a combined number of 47 CNVs, while PTX-R4, R5 and R6 had 10 (the fact that more CNVs were found in WGS samples may reflect the ease with which CNVs are called with WGS versus WES data). Potentially significant genes with CNVs were *ABCB1* (MDR1) and *ABCB4* (*MDR3*) (**Fig. 4E**) on chromosome 7 (PTX-R2a, R2b). *ABCB1* amplifications are associated with clinical resistance to PTX[49]. PTX-R4 and R5 showed structural variants on chromosome 1, and PTX-R4 show an amplification event on chromosome 17 that encompassed a variety of ABC transporters (*ABCA5, 6, 8, 9, 10*). No compelling candidate genes were found in CNVs for PTX-R6. On the other hand, inspection with IGV showed that read coverage was poor and that CNVs might not have been detected with WES data.

To confirm the importance of ABC transporters in PTX resistance, clones were treated with both PTX and verapamil, a calcium channel-blocker which can reverse ABC-transporter mediated resistance [50, 51]. We observed a complete reversal of resistance in PTX lines (**Fig. 4F**). In contrast, we observed no reversal of resistance in GEM lines (**Fig. 4G**), suggesting the resistance role of ABC-transporters is PTX-specific.

### Topotecan resistance is associated with complex alterations in *TOP1*, deletion of *WWOX* and SNVs in cytochrome p450s (*CYP1B1*)

The six TPT samples were derived from four independent selection events (TPT-R4a-c are clones from the same selection with levels of resistance ranging from 10-20 ×; **Table S2**) and all six clones were subjected to WGS together with their parent clones (WT-6 and WT-7)

For TPT-R4a-c lines (**Fig. 5A**), 268 alleles were present with AF > 0.85, but of these, only six were coding mutations and the rest were intergenic. Three of the six coding mutations were frameshift mutations (His81) with AF = 1 in *TOP1* (**Fig. 5B, S6A**), the known target of topotecan[23]. The His81 frameshift mutation, which introduces a premature stop codon, was confirmed by examining the read alignments (**Fig. S9A**) and by the absence of the full-length protein using N-terminal antibodies (**Fig. 5C**). Because there were also complex structural variants in the region (**Fig 5D, S9B**) we also sequenced the 5’ cDNA through the His81 frameshift for all three lines and as well as the parent line and confirmed the two-base deletion in the mutant as well as homozygosity in TPT-R4a-c evolved lines. We also observed a decrease in mRNA expression with TPT-R4a-b showing a statistically significant decrease in TOP1 mRNA expression, relative to TPT-WT (**Fig. S10**). It has been previously shown that a targeted RNAi suppression of Top1 produces resistance to camptothecin, a close analog of topotecan[37]. Interestingly, of the 22 TOP1 frameshift or nonsense mutations in the COSMIC tumor database, 6 were located within a 30 amino acid span (of 765 total) that includes His81 (exon 4), suggesting likely clinical relevance[52]. The probability of this distribution by chance is 9.65 ×10^−5^.

**Fig. 5.**
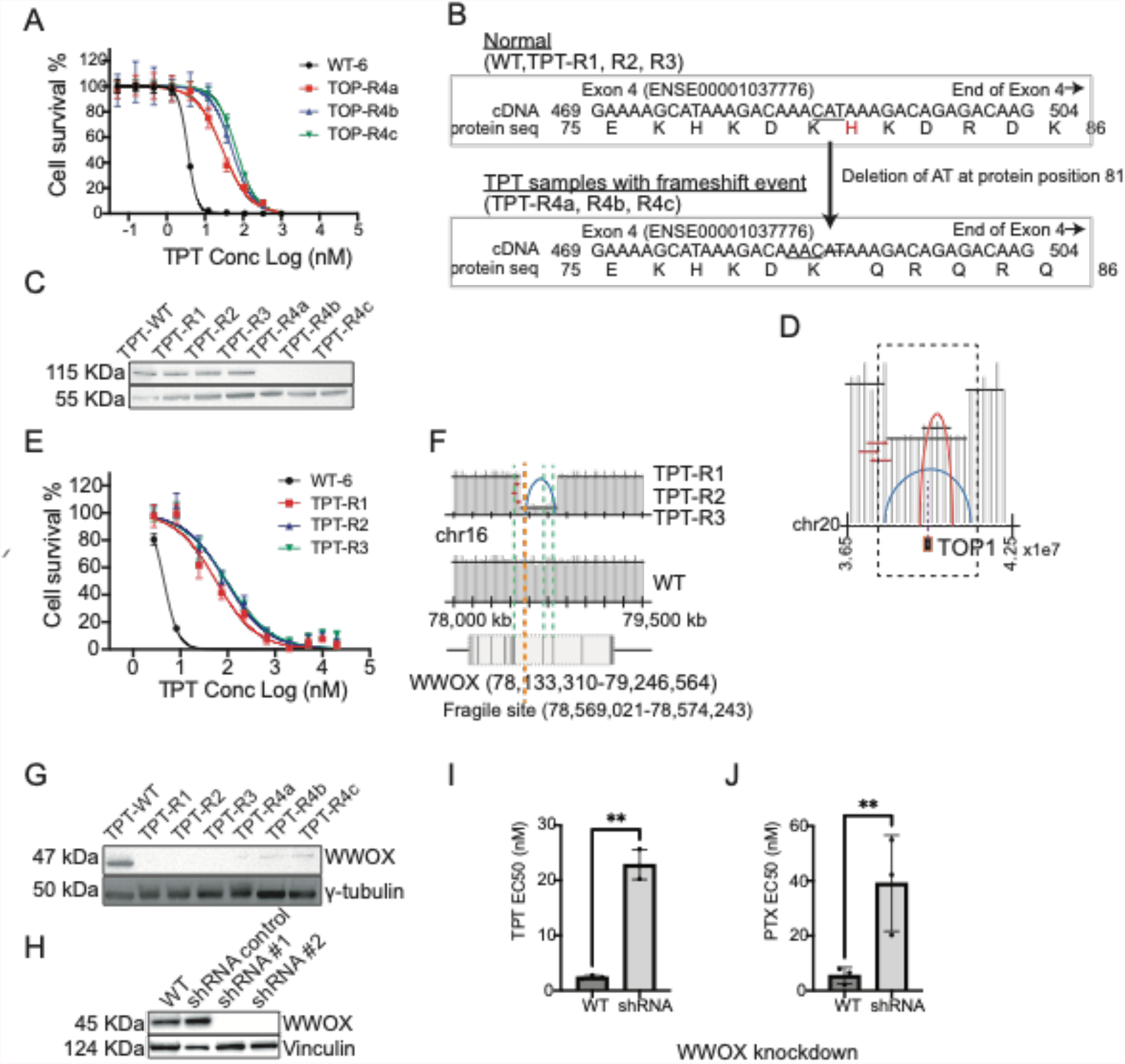
**A**. TPT EC_50_ curves for TOP-R4a,b and c. **B**. Transcript sequence (cDNA) and protein sequence for TOP1 transcript for normal (top) and TOP 4a, b and c. The figure shows only part of the cDNA (position 469-504) and protein sequence (position 75-86) for TOP1 at the affected exon (Exon 4, ENSE00001037776). The frameshift deletion of nucleotides ‘CAT’ to ‘C’ observed in TPT samples (TPT-R4a, R4b, and R4c) is predicted to give a frameshift at amino acid 81 (His, red highlight in normal). Amino acids affected by the frameshift deletion are highlighted in red. **C**. Western blot TOP1 protein depletion in evolved lines. **D**. Schematic showing complex read depth patterns around *TOP1*. ***E***. TPT EC_50_ curves for evolved TOP-R1, R2 and R3. **I**. Schematic of chr16 reads around *WWOX* for TPT-R1, R2, and R3 compared to the WT chromosome 16 parental cell line. Blue arch represents a deleted region. *WWOX* below shows the exonic (black lines) and intronic (white box) regions of the gene. The start of the deletion event is also close to a known fragile site (orange dashed line). **G**. Western blot showing WWOX protein levels in TPT resistant clones. **H**. Western blot shows downregulation of protein levels for *WWOX* in shRNA samples compared to WT and scrambled control. Barplot of the WT and shRNA knockdown cell lines for *WWOX* showing EC_50_ values for TPT (**I**) and PTX (**J**). Data is represented by mean ± s.e.m. with n=3 biological replicates and 4-8 technical replicates. ** = p < 0.05. p values determined by ratio, paired two-tailed *t* test.

No clear coding SNVs with a high allele frequency were obvious in TPT-R1, R2 and R3 but we noted multiple SNVs (Asp217Glu from TPT-R4a,b,c and Val432Leu from TPT-R1) in *CYP1B1*, which encodes a cytochrome p450 isoform. Overexpression of CYP1B1 has previously been associated with TPT resistance[50]. TPT resistant lines (TPT-R1, R2 and R3 (**Fig. 5E**) also showed large chromosomal abnormalities at *WWOX* (**Fig. 5F**) with a clear deletion of the *WWOX* gene region (chr16:78,569,166-78,792,736, exon7 and 8). WWOX bears a well-known fragile site (FRA16D) and encodes a putative oxidoreductase. The complete absence of WWOX protein was confirmed by Western in TPT-R1, 2 and 3 (**Fig. 5G**). Interestingly, lower levels of WWOX were also observed in TPT-R4a-c, which could be a consequence of other *cis* or *trans* variants in this cell line and might also contribute to this level of resistance. Knockdown of *WWOX* by shRNA resulted in marked resistance to TPT (**Fig. 5H, I**). WWOX acts as a tumor suppressor and plays a role in apoptosis. Its disruption may prevent TPT-induced apoptosis, promoting cell survival in the presence of TPT[53]. WWOX disruption also resulted in resistance to PTX (**Fig. 5J**), and as reported by others who examined WWOX-transfected epithelial ovarian cancer cells [54].

#### Some evolved mutations are associated with more drug resistance in human cancer cell lines

We further evaluated the association of mutations in our identified genes with drug resistance in cancer cell lines, reasoning that if resistance genes were already mutated before drug exposure, the cell lines would be more resistant. The set of genes we identified had mutations in multiple cancer cell lines [55, 56]. Considering only matched alteration types, we identified cell lines with mutations in the same set of resistance genes (**Table S9**). We grouped cell lines according to whether or not they had a drug-specific resistance mutation of matched type (SNV or CNV) in any resistance gene found in the HAP1 study and compared areas under the dose-response curve (AUC) between groups [24] (**Fig. S11A**). All five drugs trended toward higher dose response AUCs when resistance genes were already mutated indicating that they are more resistant, with differences in doxorubicin, gemcitabine and paclitaxel all showing significance after multiple test correction. This comparison is complicated by the fact that not all missense mutations are necessarily functional, thus some cell lines may be included in the mutated category that do not actually have altered protein function. Additionally, mutations and karyotypic abnormalities affecting other genes could also contribute to resistance in each cell line, thus cell lines lacking mutations in certain genes can still be resistant. We therefore attempted to further compare cell lines carrying mutations based on the predicted functional consequences of mutations in the context of specific gene-drug pairs. For some individual pairs, predicted loss of function (LOF) variants tended to have higher dose-response AUCs than variants predicted to have weaker effects on protein activity, in particular, SPG7 and SLC13A4 for doxorubicin, WDR33 for etoposide, and CYP1B1 for topotecan (**Fig. S11B**). Only CYP1B1 met statistical significance at a 0.1 Type 1 error rate.

## DISCUSSION and CONCLUSIONS

Here, we show for the first time that *in vitro* evolution and whole genome analysis (IVIEWGA) can readily lead to the identification of drug resistance mechanisms in human cells. Our results show *in vitro* resistance acquisition and provide a framework for the determination of chemotherapy resistance alleles that may arise in patients.

Our work using IVIEWGA in pathogens (see [55] for a review) guided our pipeline development: We focused on protein coding alterations that arose in association with a single treatment condition, that were nonsynonymous, occurring repeatedly and were high allele frequency. We also removed alleles for genes that are known to mutate frequently, like odorant receptors. Overall, our results are similar to what we have observed in eukaryotic pathogens with a mix of CNVs and single nucleotide variants giving rise to resistance.

Because of the substantially greater costs associated with WGS, here we did evaluate both WES and WGS sequencing methods. Despite a higher likelihood of discovering all changes by WGS, the disadvantage of WGS is cost and computational time. While human WES data can be analyzed on a laptop, human WGS data files are large and difficult to handle, computationally. It has been estimated, considering computational time, that a human genome costs upwards of $25,000 to fully sequence [56].

The biggest disadvantage of using WES is that CNVs will be harder to call. This is partly for statistical reasons with many reads that support CNV calls located outside of coding regions in WGS samples. In addition, if one sequences over the exact location of the recombination event (or the start or end of the deletion) one can obtain additional support for location calls via split read analysis of paired-end libraries. In addition, one can extract the sequence of the short read and reconstruct the exact recombination breakpoint, as shown in **Fig. S3**. This would not be feasible with whole exome sequencing. Recently it was shown that CNV detection tools perform poorly on WES cancer genome samples. Comparative analysis showed a low consensus in CNV calling tools with moderate sensitivity (∼50% - ∼80%), fair specificity (∼70% - ∼94%) and poor FDRs (∼27% - ∼60%). Also, using simulated data these authors observed that increasing the coverage more than 10× in exonic regions did not improve the detection power [57]. Of course, detecting CNVs is likely to be more challenging in diploid genomes, than haploid genomes. In support of this, we were able to identify and validate the RRM1 amplification event in GEM-R4, 5 and 6, which were only subjected to WES. In addition, in yeast, it appears CNVs are much less important than SNVs in driving drug resistance as well: In a more comprehensive *in vitro* evolution study in yeast with 80 different and 355 whole genome sequences we observed only 24 CNVs, including apparent aneuploidy (11 times, occurring in 10 clones) and small, intrachromosomal amplifications (13 times, occurring in 13 clones) in our set of 355 whole genome sequences [58].

A lesser disadvantage of WES, is the rare possibility that resistance is conferred by an intergenic mutation, which would be missed by WES data. Our work in other organisms has shown that almost all resistance conferring SNVs or small indels are nonsynonymous changes that would be detected by both WES and WGS. In the aforementioned yeast study, 271 mutations of the 1405 detected mutations in the 355 evolved lines were intergenic. Of these, only five were directly upstream or downstream of one of the 137 genes that were repeated identified in the study, In contrast to coding mutations, most intergenic mutations lacked any statistical support suggesting relevance and were likely to be background mutations [58]. Despite the lower probability that intergenic or other noncoding mutations may have functional effect, we recognize that there are examples from the literature where intergenic mutations have contributed to drug resistance. Non-coding RNAs such as *EGRF-AS1* and activating cis elements such as enhancers have previously been implicated in evasion of drug response[59-61]. The intergenic mutations with high allele frequency are present in our provided datasets and provide opportunity for reanalysis or for querying by those interested in a specific noncoding RNA or enhancer. It is feasible that even synonymous mutations could confer resistance if they altered the rate of protein folding.

A limitation of our HAP1 study, as presented, and is contrast to our work in other species, is that despite some level of repetition, we seldom achieved strong statistical confidence by just performing selections and sequencing. This may not be unexpected. Evolution is, unfortunately, a relatively stochastic process even when working with the exact same starting clone. In the yeast study [58] we only obtained the same allelic change in the same critical drug resistance gene a few times despite >3 repetitions per each of the 80 compounds. For example, two independent selections with hectochlorin both yielded an Arg116Lys in ACT1, the target of hectochlorin [58]. Similarly, a Leu671Phe change in YRM1 was observed 5 times for 4 different compounds.

Another disadvantage of using human cells is the challenge of validation of SNVs; we were not able to engineer any SNVs into HAP1 cells to demonstrate their importance. On the other hand, with the statistical confidence that comes from identifying the same gene repeatedly, CRISPR-*Cas9* validation becomes less important. In the same yeast study described above, YRM1, a gene encoding a transcription factor involved in drug resistance in yeast was independently identified 52 times with 27 different alleles. The likelihood of 355 selections yielding the same gene by chance is roughly 3.53 × 10^−116^. This enrichment analysis becomes an attractive method for teasing apart driver and passenger mutations and may become possible with more repetitions despite the larger genome size of HAP1 cells. However, performing enough repetitions to achieve statistical confidence would require substantial resources with WGS. even with a thousand-dollar human genome. WES is thus likely to be more useful.

While HAP1 cells may not be considered a perfect model for human cancer biology, for the purposes of target identification, they are likely very useful. As with pathogens, our use of well-studied drugs, largely uncovered genes that were mostly already well known to confer resistance such as RRM1 [62], [41], DCK [63, 64], TOP2A [37]. and TOP1[37] in a variety of different cancer cell lines. Although it was initially argued that the *in vitro* evolution system might be artificial, in malaria parasites it has been used to discover or rediscover most, if not all (to our knowledge), clinically relevant drug resistance genes including the chloroquine resistance transporter[5], the artemisinin resistance gene, *Pfkelch13*[65], and well-known ABC transporters[5].

Despite questions about how much they mimic human cells, the value of using haploid cell lines is evident from our allele frequency data. If our lines had been diploid and we would have needed to consider allele frequency data of up to 0.4. There are 205 missense mutations with an AF of >0.4, making pinpointing the causative allele much more difficult without candidate genes or without many repetitions. Although *in vitro* evolution has been used repeatedly for discovering the mechanism of action of completely uncharacterized compound in malaria parasites (reviewed in [55]), there are fewer examples of *in vitro* evolution being used for de novo target discovery in diploid eukaryotic pathogens. Although there are some examples in trypanosomes [66-68], some hypothesis about the mechanism of action was already present before evolution studies were attempted. Despite this, low allele frequency data should not necessarily be discarded. There have been examples from haploid Plasmodium where a resistance-conferring allele was located within an amplification region and thus showed an AF < 0.5.

Although the HAP1 cells could be considered unnatural, it is likely that similar evolution experiments in other types of human cells will largely give the same genes. This is because conservation of drug targets and drug resistance mechanisms across phyla is often observed, although a given compound or inhibitor may show differences in selectivity and specificity. Resistance to topotecan/camptothecan in yeast is also provided by mutations in Top1[69]. Recent IVIEWGA studies in yeast also identified Top1 as the target in yeast [58]. Evolution studies with cladosporin in yeast and plasmodium both give the same resistance mechanism for cladosporin, lysyl tRNA synthetase [70].

Our studies were not meant to study the process of evolution. Within the field of laboratory-based evolution, there are two broad areas of study. The first are those that fall under the heading of “experimental evolution” and which try to mimic evolution in natural conditions. Here, growth rates are often recorded and experimental conditions may be varied in a controlled manner (carbon sources, temperature, etc). Such studies include long term studies of *E. coli* or other bacteria(reviewed in [71]) and have also been performed with small molecules [8],[72, 73], primarily with known mechanism of action. Alternatively, there are also studies in which evolution has been used as a tool to discover targets and resistance genes for therapeutic purposes [7, 74] [55]. In many cases [74, 75], although not in all cases the term “in vitro evolution” is used instead of “experimental evolution.” Based on our results here, resistance readily emerges in HAP1 cells but more work will need to be done to determine if this is because of the compounds that were used. Here we used *in vitro* evolution (versus experimental evolution) to select for mutant lines that could withstand treatment with the selected drugs. Although it may be possible to use HAP1 cells for experimental evolution, at present sequencing costs are so high that whole genome studies with whole genome analysis are not practical but this may change in the future. Questions that might be investigated include the fitness of different mutations, reproducibility of the process, impact of the starting clone, carbon sources or growth rate and whether one resistance mechanism predominates or if a variety are found.

Finally, it is important to keep in mind that the compounds examined here are not modern cancer therapies and while still used clinically, they are imperfect. Newer molecules include bortezomib, a small molecule proteasome inhibitor, imatinib, a small molecule tyrosine kinase inhibitor or seliciclib, small molecule cyclin-dependent kinase inhibitor or even small molecule cancer immunotherapies. We anticipate mutations in the drug’s targets will be identified sometimes, as is observed in microbes. In fact, unbiased IVIEWGA studies with bortezomid in *P. falciparum* have demonstrated mutations in the proteosome subunit, Pf20S β5, [76] confer resistance, and similar resistance-conferring mutations have been discovered after using in vitro evolution in human cells, although whole genome sequencing was not performed and the mutations were identified using a candidate gene approach [77]. On the other hand, targeted therapies are less likely to work against HAP1 cells, as shown here for imatinib (**Fig. S1**), most likely because HAP1 cells do not harbor the appropriate sensitizing mutations (e.g. the BRC-Abl for imatinib or BRAF/EGFR mutations for vemurafenib, gefitinib or erlotinib, respectively [78]). Alternatively, the HAP1 cells may be intrinsically resistant because they harbor other resistance conferring mutations. Nevertheless, if they can be used or engineered to sensitivity, predicting resistance mechanisms for new drugs in clinical development, as well as for new drug combinations and may lead to better classes of drugs for chemotherapy.

## MATERIALS AND METHODS

### Compounds

All chemotherapeutic agents used in this study were obtained from Sigma-Aldrich, dissolved in DMSO at 10mM concentration and stored at -20°C.

### Cell cultures

The human chronic myelogenous leukemia cell line, HAP1, was purchased as authenticated at passage 7 from Horizon Discovery and cultured in tissue culture dishes (Genesee Scientific, Cat# 25-202) as a monolayer at 37°C in a humidified atmosphere with 5% CO_2_ using Iscove’s Modified Dulbecco’s Medium (IMDM) (Life Technologies, CA) supplemented with 10% fetal bovine serum (FBS), 0.29mg/mL L-glutamine, 25mM HEPES, 100U/mL Penicillin and 100µg/mL Streptomycin (1% pencillin/streptomycin). Monoclonal and polyclonal stocks were made and stored in IMDM + 10% DMSO in liquid nitrogen.

### *In vitro* evolution of resistant HAP1 clones

Prior to selection, an aliquot of the parental line was stocked as a reference for subsequent whole genome sequencing analysis. Three independent clones of HAP1 cells were cultured in tissue culture dishes exposed to increasing sublethal concentrations of each chemotherapeutic agent at a starting concentration previously determined by the EC_50_ value for around 7-30 weeks depending on the drug, its speed of action and the method used as two methods were considered: high-pressure intermittent selection method and a step-wise selection method. For high pressure selection, cells were treated at a concentration 3-10 × EC_50_ value until more than 95% of the cells died. Then treatment was removed and cells were allowed to recover. After reaching around 60% semi-confluence, treatment was reinstalled and EC_50_ value monitored. For step-wise selection method, drug concentration used was at the EC_50_ which implied reduced growth rate of approximately 50% and drug pressure was increased in intervals of around 5-10% keeping growth inhibition around 50%. Once the EC_50_ values of the resistant lines were at least 5 times greater than the one used as control, cells were again cloned by limiting dilution and further reconfirmed for drug resistance and subsequent DNA extraction for whole genome sequencing analysis.

### Dose-response assay by EC_50_ determination and bioluminescence quantification

Drug sensitivity and cell viability were assessed by a bioluminescence measurement as follows: twenty-four hours prior to addition of the drugs, 2 ×10^4^ cells/well for every replicate were seeded in a 96-well plate. Once attached, media was removed and 10 different concentrations of drug were added in serial dilutions 1:3 with a starting concentration of 10µM or one of which the EC_50_ value of the replicates fell within an intermediate drug concentration. When drug-resistant lines were co-treated in combination with verapamil, a fixed concentration of verapamil (10µM) was added to every concentration of the drug. After a 48-hour incubation period at 37°C and 5% CO_2_ with the drug, cells were treated with CellTiterGlo (Promega) reagent (diluted 1:2 with deionized water) for quantification of HAP1 cell viability. Immediately after addition of the luminescence reagent, luminescence was measured using the Synergy HT Microplate Reader Siafrtd (BioTek). The data was normalized to 100% cell survival and 100% cell death and EC_50_ values were obtained using the average normalized luminescence intensity of 8 wells per concentration and a non-linear variable slope four-parameter regression curve fitting model in Prism 8 (GraphPad Software Inc.). Unless otherwise noted, dose response experiments consisted of 4-8 technical replicates and 3 biological replicates.

### Isolation of total DNA from drug resistant lines

Genomic DNA (gDNA) was extracted from drug-specific resistant cell lines together with their isogenic parental lines using the DNeasy^®^ Blood & Tissue Kit (Qiagen) following the manufacturer’s instructions. Samples were assessed for quantity with the Qubit^™^ dsDNA BR Assay Kit (Life Technologies, Carlsbad, CA, USA). All samples (>2.0µg, >50ng/µL, >20µL) were prepared for quality control by testing gDNA degradation or potential contamination using agarose gel electrophoresis (1% Agarose, TAE, ∼100 Voltage). Then gDNA concentration was again measured using the Qubit^®^ DNA Assay Kit with the Qubit^®^ 2.0 Fluorometer (Life Technologies). Finally, fragment distribution of the gDNA library was measured using the DNA 1000 Assay Kit with the Agilent Bioanalyzer 2100 system (Agilent Technologies, Santa Clara, CA, USA). DNA libraries were sequenced with 150 base pair (bp) paired single end reads on an Illumina HiSeq 4000 (PE150).

### Genome Sequencing and Data Analysis

The quality of the raw FASTQ files was checked with FastQC (http://www.bioinformatics.babraham.ac.uk/projects/fastqc/). Whole genome sequencing (WGS) reads were mapped to GRCh37 (hg19) using BWA (v.0.7.17), specifically with the hs37d5 reference genome from 1000 Genomes project (Phase II). Whole exome sequencing (WES) samples were captured using Agilent SureSelect Human All Exon V6 (58 M), and the reads were also mapped to GRCh37 using BWA (v.0.7.17) with the same reference genome as WGS. Duplicate reads were removed using Picard (v.1.94); paired resistant and parent (WT) BAM files were used as input for The Genome Analysis Toolkit (GATK, v3.8-1). Local realignment and base quality recalibration were performed using default parameters. Somatic single nucleotide variants (SNVs) and small insertion and deletion (indels) were called using GATK MuTect2 following the state-of-the-art GATK Best Practices pipeline (https://ccbr.github.io/Pipeliner/Tools/MuTect2.html). In this project, the input to MuTect2 consisted of alignments for the parent and resistant clone in order. to call mutations with statistically significant differences in read support in the setting of resistance. Only the variants with PASS status, suggesting confident somatic mutations, were considered for further analysis.

Variant allelic fraction was determined as the fraction of reads supporting the variant allele relative to total read depth at the variant position. Minimum callable depth was set to 10 and base quality score threshold was set to 18, following the default from MuTect2. All sequences have been deposited in SRA BioProject PRJNA603390.

### Small-Variant Annotation for SNVs and Indels

Somatic variants were annotated using snpEff (v 4.3q)[79]. The annotation was performed with parameters including (1) canonical transcripts and (2) protein coding to enable identification of different functional classes of variant and their impact on protein coding genes (Table 1 showing finalized and consolidated annotations; Table S4 shows the raw annotation from snpEff and consolidated classification used in Table 1; Table S7 shows all the SNVs with their raw annotations). The snpEff sequence ontology designation was used in the filtering steps to classify variants generally as noncoding or coding (Table S4).

### Identification of Drug Specific Genes

First, we excluded all variants in non-coding regions. Second, we excluded all non-functional variants, retaining only variants with a snpEff definition of HIGH or MODERATE impact (missense, stop lost, stop gain, and structural interaction variants). Finally, we selected only the variants with high allele frequency (AF > 0.85) and genes with multiple independent amino acid changes found in the same drug as the final list of candidates. The potential candidate variants were evaluated through Integrative Genomics Viewer (IGV)[80] to ensure coverage and allele fractions of the mutation positions. The top genes for each drug were included in Table 2 and Table S8.

### Somatic Copy Number Variations (CNVs) Analysis

Copy number regions for WGS and WES were called by ControlFreeC^47^ using the default settings for WGS and WES data. Background parental clone samples for each drug served as the control. Recurrent CNV regions were defined as regions observed in more than 1 sample, but not in all of clones from the tested drugs (as these are more likely to indicate potential sequencing artifacts).

### Gene knockdowns

shRNAs targeting *TOP2A* (Cat# sc-36695-V), *DCK* (Cat# sc-60509-V), *SLCO3A1* (Cat# sc-62713-V), *SLC13A4* (Cat# sc-89760-V), *KLF-1* (Cat# sc-37831-V), *WWOX* (Cat# sc-44193-V), *WDR33* (Cat# sc-94735-V) and the non-coding control (Cat# sc-108080) were obtained in pLKO.1-Puro from Santa Cruz Biotechnology. *RRM1* (clone ID NM_001033.2-476s1c1) and *CYP1B1* (clone ID NM_000104.2-1176s1c1) were obtained in pLKO.1-Puro-CMV-tGFP from Sigma Aldrich.

Gene expression was knocked down using either a shRNA pool (Santa Cruz Biotechnology) containing between three and five expression constructs each encoding target-specific 19-25 shRNAs or a single shRNA (Sigma Aldrich). HAP1 cells were plated at 120,000 cells per well (∼40% confluency) in a 24-well plate 24 hours prior to viral transduction. On the day of transduction, complete media was replaced with serum-free media and 7µg/mL Polybrene^®^ (Cat# sc-134220) and virus was added to the cells at a multiplicity of infection of 0.5 and cells were incubated overnight at 37°C. The following day, media was replaced with complete media without Polybrene and cells were incubated at 37°C overnight. Cells were then split 1:3 and incubated for 24 hours more and finally stable clones expressing the shRNA were selected using complete media with 2µg/mL puromycin. After 7 days of selection with puromycin, knockdown efficiency was confirmed by western blot. Cells transduced with shRNAs containing fluorescent tags, were trypsinized (TrypLE^™^ Express; Cat# 12605-010, Gibco) after puromycin selection, washed twice with DPBS (1X) (Gibco) and sorted by flow cytometry.

### Knockout of USP47

*USP47* was knocked out (Cat# HSPD0000092816) using a single plasmid CRISPR-*Cas9* system, using as lentivirus backbone the LV01 U6-gRNA:ef1a-puro-2A-Cas9-2A-tGFP targeting *USP47* (Sigma Aldrich). The target sequence (5’-3’) was CAATGGGGCTTCTACTAGG. Transduction was as described above. HAP1 cells were plated at 120,000 cells per well (∼40% confluency) in a 24-well plate 24 hours prior to viral transduction. On the day of transduction, complete media was replaced with serum-free media and 7µg/mL Polybrene^®^ (Cat# sc-134220), virus was added to the cells at a multiplicity of infection of 0.5 and cells were incubated overnight at 37°C. The following day, media was replaced with complete media without Polybrene and cells were incubated at 37°C overnight. Cells were then split 1:3 for 24 hours more and stable clones expressing the CRISPR-*Cas9* sequence were selected using complete media with 2µg/Ml puromycin. After 14 days of selection with puromycin and propagation as required, cells were trypsinized (TrypLE^™^ Express; Cat# 12605-010, Gibco), washed twice with DPBS (1X) (Gibco) and sorted by flow cytometry using the GFP fluorochrome which is expressed with Cas9. GFP positive cells were plated at an average density of 0.5 cells per well in a 96-well plate (previously treated with poly-L-Lysine (Sigma #P4707-50ml) to improve cell adhesion) in presence of 2µg/mL puromycin (limiting dilution cloning). Cell growth was monitored via microscopy during 25 days to select those wells which were observed to contain single colonies and *USP47* knockout was confirmed in those monoclonal HAP1 cell lines first via PCR and then reconfirmed by western blot using the *USP47* rabbit polyclonal antibody (Abcam, Cat# ab97835).

### Immunoblotting

HAP1 cells (at least 5 ×10^6^) were trypsinized, washed twice with cold 1 × DPBS and then lysed in 500µL Pierce^™^ RIPA Buffer (Thermo Scientific) containing 1:100 protease inhibitor (Halt^™^ Protease & Phosphatase Inhibitor Cocktail, Thermo Scientific) and 1:100 0.5M EDTA Solution (Thermo Scientific). Total protein concentration was measured using the DC Protein Assay (Bio-Rad). Equal amounts of proteins were resolved by SDS-PAGE and transferred to nitrocellulose membranes (Bio-Rad #1704271), blocked in PBS with 5% (w/v) Blotting-Grade Blocker (Bio-Rad #170-6404) and 0.1% (v/v) Tween20 for 1h and probed. As secondary antibodies, HRP-linked anti-mouse or anti-rabbit (Cell Signaling Technology) were used and the HRP signal was visualized with SuperSignal^®^West Pico Chemiluminescent Substrate (Thermo Scientific #34080) using Syngene G-Box imager. Protein enrichment was calculated relative to *vinculin, γ-tubulin* or *β-actin*. Primary antibodies are listed below. Full size western blots are shown in **Fig. S12**.

### Antibodies

*TOP2A* (Sigma #SAB4502997), *USP47* (Abcam #ab97835), *WDR33* (Abcam #ab72115), *DCK* (Abcam #ab151966), *β-actin* (Cell Signaling #3700S), *γ-tubulin* (Cell Signaling #4285S), *Vinculin* (Invitrogen #700062), *SLC13A4/SUT-1* (Abcam #ab236619), *WWOX* (Abcam #ab137726), *EKLF/KLF-1* (Abcam #175372), *SLCO3A1/OATP-A* (Santa Cruz #sc-365007), *TOP1* (Proteintech #20705-1-AP), CRISPR-*Cas9* (Sigma #SAB4200701), *RRM1* (Abcam #ab133690), *CYP1B1* (Abcam #ab137562), *SPG7* (Sigma #SAB1406470 and Abcam #ab96213), goat anti-mouse (Invitrogen #G21040), goat anti-rabbit (Invitrogen #G21234).

### RNA isolation, RT-PCR analysis and sequencing of TOP1 (*His81*)

TPT-resistant cells and TPT-WT (1 ×10^6^ cells) were dissociated from plates by the addition of 2mL of TrypLE (Cat #12605-010, Gibco), washed and total RNA was isolated and purified using a Qiagen RNeasy® Mini Kit (Cat #74104, Qiagen) according to manufacturer’s instructions. cDNA was synthesized from 1µg of total RNA using the Superscript^™^ First-Strand Synthesis System for RT-PCR Kit (Invitrogen #11904-018) and random hexamers. The primers used to amplify the region containing *His81* were FWD: GATCGAGAACACCGGCAC and REV: TCAGCATCATCCTCATCTCGAG. DNA from PCR product was extracted, using the QIAquick® Gel Extraction Kit (Qiagen #28706) following the manufacturer’s instruction, measured using the Qubit^®^ DNA Assay Kit with the Qubit^®^ 2.0 Fluorometer (Life Technologies), and sequenced. The cDNA was sent to Eton Biosciences for Sanger sequencing. Quantification of TOP1 expression was performed using PerfeCTa^®^ Sybr Green Fast Mix (Quanta #95072-250) the following primers: FWD: CGAATCATGCCCGAGGATATAA; REV: CCAGGAAACCAGCCAAGTAA, following the manufacturer’s instruction.

### GDSC analysis methods

Mutations and copy number data for cancer cell lines were obtained from the DepMap 2021 quarter 1 release via the DepMap portal (https://depmap.org/portal/download/) on 02/01/2021. Copy number alterations in genes were determined by filtering for a log_2_(copy number + 1) greater in absolute value than 1.5. Cell lines were first grouped according to whether they had a mutation or copy number alteration that matched any of those found in Table 2. EC_50_ and dose-response area under the curve (AUC) data were obtained from the GDSC 8.3 Release (June 2020). Dose-response AUC distributions for doxorubicin, etoposide, gemcitabine, paclitaxel, and topotecan were compared between the cell lines with or without a mutation using the Mann-Whitney U test (**Fig. S11A**). P-values were corrected for multiple testing using the Benjamini-Hochberg (BH) method[81]. All cell lines with SNVs in the genes listed in Table 2 were then grouped based on functional predictions by the Variant Effect Scoring Tool (VEST4.0)[82]. Cell lines with a mutation that had a VEST score > 0.8 were labeled as “Likely LOF” cell lines, whereas cell lines with mutations that scored <= 0.8 or that were silent were labeled as “No Likely LOF Mutation” cell lines. As most predicted functional mutations result in loss of function, we assumed this was the likely consequence, though it is possible that some high scoring mutations could in fact be gain of function. Dose-response AUC distributions for these groups were then compared for each gene using the Mann-Whitney U test (**Fig. S11B**) and p-values corrected by the BH procedure.

## Supporting information

File S3. Sequencing characteristics

S5. Summary of mutation types

Table S6 SNVs

Table S8 CNVs

## ABBREVIATIONS

AF: allele frequency
CNV: Copy Number Variation
NGS: Next Generation Sequencing
WES: Whole Exome Sequencing
WGS: Whole Genome Sequencing
CML: Chronic Myelogenous Leukemia
IVIEWGA: *In Vitro* Evolution and Whole Genome Analysis
SNV: Single Nucleotide Variant
CNV: Copy Number Variation
TF: transcription factor
DOX: Doxorubicin
GEM: Gemcitabine
ETP: Etoposide
PTX: Paclitaxel
TPT: Topotecan
AML: Acute Myeloid Leukemia
TKIs: Tyrosine Kinase Inhibitors
MDR: Multi-Drug Resistance
gDNA: genomic DNA

## Acknowledgments

This work was supported by the National Institute of Health (NIH) to EAW, HC, and TI (GM085764), the San Diego Center for Systems Biology and UC San Diego Health Science fellowship to JCJ, and NIH National Library of Medicine Training Grant T15LM011271 to MD. EAW is also supported by grants from the Bill and Melinda Gates Foundation and the Medicines for Malaria Venture. The authors declare no conflicts of interest.

## Author contributions

JCJ performed selections, validation experiments and wrote the manuscript. MD, AK performed all computational analyses, and assembled figures and tables and wrote the manuscript. GF assisted with CRISPR-*Cas9* experiments. KC performed RNA extraction and RT-qPCR experiments. AK performed sequence analysis. HC provided advice and management and wrote the manuscript. EAW performed data analysis, provided advice, obtained funding and wrote the manuscript. TI obtained funding and provided advice.

## References

1. Shalem O, Sanjana NE, Hartenian E, Shi X, Scott DA, Mikkelson T, Heckl D, Ebert BL, Root DE, Doench JG, Zhang F: Genome-scale CRISPR-Cas9 knockout screening in human cells. Science 2014, 343:84–87.

2. Kanarek N, Keys HR, Cantor JR, Lewis CA, Chan SH, Kunchok T, Abu-Remaileh M, Freinkman E, Schweitzer LD, Sabatini DM: Histidine catabolism is a major determinant of methotrexate sensitivity. Nature 2018, 559:632–636.

3. Wang T, Wei JJ, Sabatini DM, Lander ES: Genetic screens in human cells using the CRISPR-Cas9 system. Science 2014, 343:80–84.

4. Antonova-Koch Y, Meister S, Abraham M, Luth MR, Ottilie S, Lukens AK, Sakata-Kato T, Vanaerschot M, Owen E, Jado JC, et al: Open-source discovery of chemical leads for next-generation chemoprotective antimalarials. Science 2018, 362.

5. Cowell AN, Istvan ES, Lukens AK, Gomez-Lorenzo MG, Vanaerschot M, Sakata-Kato T, Flannery EL, Magistrado P, Owen E, Abraham M, et al: Mapping the malaria parasite druggable genome by using in vitro evolution and chemogenomics. Science 2018, 359:191–199.

6. Ottilie S, Goldgof GM, Calvet CM, Jennings GK, LaMonte G, Schenken J, Vigil E, Kumar P, McCall LI, Lopes ES, et al: Rapid Chagas Disease Drug Target Discovery Using Directed Evolution in Drug-Sensitive Yeast. ACS Chem Biol 2017, 12:422–434.

7. Andries K, Verhasselt P, Guillemont J, Gohlmann HW, Neefs JM, Winkler H, Van Gestel J, Timmerman P, Zhu M, Lee E, et al: A diarylquinoline drug active on the ATP synthase of Mycobacterium tuberculosis. Science 2005, 307:223–227.

8. Santos-Lopez A, Marshall CW, Scribner MR, Snyder DJ, Cooper VS: Evolutionary pathways to antibiotic resistance are dependent upon environmental structure and bacterial lifestyle. Elife 2019, 8.

9. Burckstummer T, Banning C, Hainzl P, Schobesberger R, Kerzendorfer C, Pauler FM, Chen D, Them N, Schischlik F, Rebsamen M, et al: A reversible gene trap collection empowers haploid genetics in human cells. Nat Methods 2013, 10:965–971.

10. Carette JE, Raaben M, Wong AC, Herbert AS, Obernosterer G, Mulherkar N, Kuehne AI, Kranzusch PJ, Griffin AM, Ruthel G, et al: Ebola virus entry requires the cholesterol transporter Niemann-Pick C1. Nature 2011, 477:340–343.

11. Gerhards NM, Blomen VA, Mutlu M, Nieuwenhuis J, Howald D, Guyader C, Jonkers J, Brummelkamp TR, Rottenberg S: Haploid genetic screens identify genetic vulnerabilities to microtubule-targeting agents. Mol Oncol 2018, 12:953–971.

12. Findlay GM, Daza RM, Martin B, Zhang MD, Leith AP, Gasperini M, Janizek JD, Huang X, Starita LM, Shendure J: Accurate classification of BRCA1 variants with saturation genome editing. Nature 2018, 562:217–222.

13. Behnisch-Cornwell S, Wolff L, Bednarski PJ: The Effect of Glutathione Peroxidase-1 Knockout on Anticancer Drug Sensitivities and Reactive Oxygen Species in Haploid HAP-1 Cells. Antioxidants (Basel) 2020, 9.

14. Smits AH, Ziebell F, Joberty G, Zinn N, Mueller WF, Clauder-Münster S, Eberhard D, Fälth Savitski M, Grandi P, Jakob P, et al: Biological plasticity rescues target activity in CRISPR knock outs. Nat Methods 2019, 16:1087–1093.

15. Essletzbichler P, Konopka T, Santoro F, Chen D, Gapp BV, Kralovics R, Brummelkamp TR, Nijman SM, Bürckstümmer T: Megabase-scale deletion using CRISPR/Cas9 to generate a fully haploid human cell line. Genome Res 2014, 24:2059–2065.

16. Mandell JB, Lu F, Fisch M, Beumer JH, Guo J, Watters RJ, Weiss KR: Combination Therapy with Disulfiram, Copper, and Doxorubicin for Osteosarcoma: In Vitro Support for a Novel Drug Repurposing Strategy. Sarcoma 2019, 2019:1320201.

17. Jiang C, Jiang L, Li Q, Liu X, Zhang T, Yang G, Zhang C, Wang N, Sun X, Jiang L: Pyrroloquinoline quinine ameliorates doxorubicin-induced autophagy-dependent apoptosis via lysosomal-mitochondrial axis in vascular endothelial cells. Toxicology 2019, 425:152238.

18. Drenberg CD, Shelat A, Dang J, Cotton A, Orwick SJ, Li M, Jeon JY, Fu Q, Buelow DR, Pioso M, et al: A high-throughput screen indicates gemcitabine and JAK inhibitors may be useful for treating pediatric AML. Nat Commun 2019, 10:2189.

19. Alvarellos ML, Lamba J, Sangkuhl K, Thorn CF, Wang L, Klein DJ, Altman RB, Klein TE: PharmGKB summary: gemcitabine pathway. Pharmacogenet Genomics 2014, 24:564–574.

20. Mini E, Nobili S, Caciagli B, Landini I, Mazzei T: Cellular pharmacology of gemcitabine. Ann Oncol 2006, 17 Suppl 5:v7–12.

21. Pommier Y, Leo E, Zhang H, Marchand C: DNA topoisomerases and their poisoning by anticancer and antibacterial drugs. Chem Biol 2010, 17:421–433.

22. Hayashi MT, Karlseder J: DNA damage associated with mitosis and cytokinesis failure. Oncogene 2013, 32:4593–4601.

23. Rasheed ZA, Rubin EH: Mechanisms of resistance to topoisomerase I-targeting drugs. Oncogene 2003, 22:7296–7304.

24. Yang W, Soares J, Greninger P, Edelman EJ, Lightfoot H, Forbes S, Bindal N, Beare D, Smith JA, Thompson IR, et al: Genomics of Drug Sensitivity in Cancer (GDSC): a resource for therapeutic biomarker discovery in cancer cells. Nucleic Acids Res 2013, 41:D955–961.

25. Iorio F, Knijnenburg TA, Vis DJ, Bignell GR, Menden MP, Schubert M, Aben N, Goncalves E, Barthorpe S, Lightfoot H, et al: A Landscape of Pharmacogenomic Interactions in Cancer. Cell 2016, 166:740–754.

26. Corey VC, Lukens AK, Istvan ES, Lee MCS, Franco V, Magistrado P, Coburn-Flynn O, Sakata-Kato T, Fuchs O, Gnadig NF, et al: A broad analysis of resistance development in the malaria parasite. Nat Commun 2016, 7:11901.

27. Christowitz C, Davis T, Isaacs A, van Niekerk G, Hattingh S, Engelbrecht AM: Mechanisms of doxorubicin-induced drug resistance and drug resistant tumour growth in a murine breast tumour model. BMC Cancer 2019, 19:757.

28. Amrutkar M, Gladhaug IP: Pancreatic Cancer Chemoresistance to Gemcitabine. Cancers (Basel) 2017, 9.

29. Pan P, Li Y, Yu H, Sun H, Hou T: Molecular principle of topotecan resistance by topoisomerase I mutations through molecular modeling approaches. J Chem Inf Model 2013, 53:997–1006.

30. Andersson BS, Collins VP, Kurzrock R, Larkin DW, Childs C, Ost A, Cork A, Trujillo JM, Freireich EJ, Siciliano MJ, et al.: KBM-7, a human myeloid leukemia cell line with double Philadelphia chromosomes lacking normal c-ABL and BCR transcripts. Leukemia 1995, 9:2100–2108.

31. Brammeld JS, Petljak M, Martincorena I, Williams SP, Alonso LG, Dalmases A, Bellosillo B, Robles-Espinoza CD, Price S, Barthorpe S, et al: Genome-wide chemical mutagenesis screens allow unbiased saturation of the cancer genome and identification of drug resistance mutations. Genome Res 2017, 27:613–625.

32. Boeva V, Popova T, Bleakley K, Chiche P, Cappo J, Schleiermacher G, Janoueix-Lerosey I, Delattre O, Barillot E: Control-FREEC: a tool for assessing copy number and allelic content using next-generation sequencing data. Bioinformatics 2012, 28:423–425.

33. Jeon KH, Yu HV, Kwon Y: Hyperactivated m-calpain affects acquisition of doxorubicin resistance in breast cancer cells. Biochim Biophys Acta Gen Subj 2018, 1862:1126–1133.

34. Turner JG, Dawson JL, Grant S, Shain KH, Dalton WS, Dai Y, Meads M, Baz R, Kauffman M, Shacham S, Sullivan DM: Treatment of acquired drug resistance in multiple myeloma by combination therapy with XPO1 and topoisomerase II inhibitors. J Hematol Oncol 2016, 9:73.

35. Ghisoni E, Maggiorotto F, Borella F, Mittica G, Genta S, Giannone G, Katsaros D, Sciarrillo A, Ferrero A, Sarotto I, et al: TOP2A as marker of response to pegylated lyposomal doxorubicin (PLD) in epithelial ovarian cancers. J Ovarian Res 2019, 12:17.

36. Wendorff TJ, Schmidt BH, Heslop P, Austin CA, Berger JM: The structure of DNA-bound human topoisomerase II alpha: conformational mechanisms for coordinating inter-subunit interactions with DNA cleavage. J Mol Biol 2012, 424:109–124.

37. Burgess DJ, Doles J, Zender L, Xue W, Ma B, McCombie WR, Hannon GJ, Lowe SW, Hemann MT: Topoisomerase levels determine chemotherapy response in vitro and in vivo. Proc Natl Acad Sci U S A 2008, 105:9053–9058.

38. Lim MY, LaMonte G, Lee MCS, Reimer C, Tan BH, Corey V, Tjahjadi BF, Chua A, Nachon M, Wintjens R, et al: UDP-galactose and acetyl-CoA transporters as Plasmodium multidrug resistance genes. Nat Microbiol 2016, 1:16166.

39. Mansoori B, Mohammadi A, Davudian S, Shirjang S, Baradaran B: The Different Mechanisms of Cancer Drug Resistance: A Brief Review. Adv Pharm Bull 2017, 7:339–348.

40. Mandt REK, Lafuente-Monasterio MJ, Sakata-Kato T, Luth MR, Segura D, Pablos-Tanarro A, Viera S, Magan N, Ottilie S, Winzeler EA, et al: In vitro selection predicts malaria parasite resistance to dihydroorotate dehydrogenase inhibitors in a mouse infection model. Sci Transl Med 2019, 11.

41. Bepler G, Kusmartseva I, Sharma S, Gautam A, Cantor A, Sharma A, Simon G: RRM1 modulated in vitro and in vivo efficacy of gemcitabine and platinum in non-small-cell lung cancer. J Clin Oncol 2006, 24:4731–4737.

42. Zeng C, Fan W, Zhang X: RRM1 expression is associated with the outcome of gemcitabine-based treatment of non-small cell lung cancer patients--a short report. Cell Oncol (Dordr) 2015, 38:319–325.

43. Yang Z, Fu B, Zhou L, Xu J, Hao P, Fang Z: RRM1 predicts clinical outcome of high- and intermediate-risk non-muscle-invasive bladder cancer patients treated with intravesical gemcitabine monotherapy. BMC Urol 2019, 19:69.

44. Chan SL, Huppertz I, Yao C, Weng L, Moresco JJ, Yates JR, 3rd, Ule J, Manley JL, Shi Y: CPSF30 and Wdr33 directly bind to AAUAAA in mammalian mRNA 3’ processing. Genes Dev 2014, 28:2370–2380.

45. Teloni F, Michelena J, Lezaja A, Kilic S, Ambrosi C, Menon S, Dobrovolna J, Imhof R, Janscak P, Baubec T, Altmeyer M: Efficient Pre-mRNA Cleavage Prevents Replication-Stress-Associated Genome Instability. Mol Cell 2019, 73:670–683 e612.

46. Hansen LT, Lundin C, Spang-Thomsen M, Petersen LN, Helleday T: The role of RAD51 in etoposide (VP16) resistance in small cell lung cancer. Int J Cancer 2003, 105:472–479.

47. Huang F, Mazin AV: A small molecule inhibitor of human RAD51 potentiates breast cancer cell killing by therapeutic agents in mouse xenografts. PLoS One 2014, 9:e100993.

48. Thakkar N, Lockhart AC, Lee W: Role of Organic Anion-Transporting Polypeptides (OATPs) in Cancer Therapy. Aaps j 2015, 17:535–545.

49. Vaidyanathan A, Sawers L, Gannon AL, Chakravarty P, Scott AL, Bray SE, Ferguson MJ, Smith G: ABCB1 (MDR1) induction defines a common resistance mechanism in paclitaxel-and olaparib-resistant ovarian cancer cells. Br J Cancer 2016, 115:431–441.

50. Cruz-Munoz W, Di Desidero T, Man S, Xu P, Jaramillo ML, Hashimoto K, Collins C, Banville M, O’Connor-McCourt MD, Kerbel RS: Analysis of acquired resistance to metronomic oral topotecan chemotherapy plus pazopanib after prolonged preclinical potent responsiveness in advanced ovarian cancer. Angiogenesis 2014, 17:661–673.

51. Wu CP, Calcagno AM, Ambudkar SV: Reversal of ABC drug transporter-mediated multidrug resistance in cancer cells: evaluation of current strategies. Curr Mol Pharmacol 2008, 1:93–105.

52. Forbes SA, Beare D, Boutselakis H, Bamford S, Bindal N, Tate J, Cole CG, Ward S, Dawson E, Ponting L, et al: COSMIC: somatic cancer genetics at high-resolution. Nucleic Acids Res 2017, 45:D777–d783.

53. Del Mare S, Salah Z, Aqeilan RI: WWOX: its genomics, partners, and functions. J Cell Biochem 2009, 108:737–745.

54. Janczar S, Nautiyal J, Xiao Y, Curry E, Sun M, Zanini E, Paige AJ, Gabra H: WWOX sensitises ovarian cancer cells to paclitaxel via modulation of the ER stress response. Cell Death Dis 2017, 8:e2955.

55. Luth MR, Gupta P, Ottilie S, Winzeler EA: Using in Vitro Evolution and Whole Genome Analysis To Discover Next Generation Targets for Antimalarial Drug Discovery. ACS Infect Dis 2018, 4:301–314.

56. Schwarze K, Buchanan J, Taylor JC, Wordsworth S: Are whole-exome and whole-genome sequencing approaches cost-effective? A systematic review of the literature. Genet Med 2018, 20:1122–1130.

57. Zare F, Dow M, Monteleone N, Hosny A, Nabavi S: An evaluation of copy number variation detection tools for cancer using whole exome sequencing data. BMC Bioinformatics 2017, 18:286.

58. Ottilie S, Luth MR, Hellemann E, Goldgof GM, Vigil E, Kumar P, Cheung AL, Song M, Godinez-Macias KP, Carolino K, et al: Defining the Yeast Resistome through <em>in vitro</em> Evolution and Whole Genome Sequencing. bioRxiv 2021:2021.2002.2017.430112.

59. Si W, Shen J, Zheng H, Fan W: The role and mechanisms of action of microRNAs in cancer drug resistance. Clin Epigenetics 2019, 11:25.

60. Sanchez Calle A, Kawamura Y, Yamamoto Y, Takeshita F, Ochiya T: Emerging roles of long non-coding RNA in cancer. Cancer Sci 2018, 109:2093–2100.

61. Sur I, Taipale J: The role of enhancers in cancer. Nat Rev Cancer 2016, 16:483–493.

62. Minami K, Shinsato Y, Yamamoto M, Takahashi H, Zhang S, Nishizawa Y, Tabata S, Ikeda R, Kawahara K, Tsujikawa K, et al: Ribonucleotide reductase is an effective target to overcome gemcitabine resistance in gemcitabine-resistant pancreatic cancer cells with dual resistant factors. J Pharmacol Sci 2015, 127:319–325.

63. Saiki Y, Yoshino Y, Fujimura H, Manabe T, Kudo Y, Shimada M, Mano N, Nakano T, Lee Y, Shimizu S, et al: DCK is frequently inactivated in acquired gemcitabine-resistant human cancer cells. Biochem Biophys Res Commun 2012, 421:98–104.

64. Ohhashi S, Ohuchida K, Mizumoto K, Fujita H, Egami T, Yu J, Toma H, Sadatomi S, Nagai E, Tanaka M: Down-regulation of deoxycytidine kinase enhances acquired resistance to gemcitabine in pancreatic cancer. Anticancer Res 2008, 28:2205–2212.

65. Ariey F, Witkowski B, Amaratunga C, Beghain J, Langlois AC, Khim N, Kim S, Duru V, Bouchier C, Ma L, et al: A molecular marker of artemisinin-resistant Plasmodium falciparum malaria. Nature 2014, 505:50–55.

66. Khare S, Roach SL, Barnes SW, Hoepfner D, Walker JR, Chatterjee AK, Neitz RJ, Arkin MR, McNamara CW, Ballard J, et al: Utilizing Chemical Genomics to Identify Cytochrome b as a Novel Drug Target for Chagas Disease. PLoS Pathog 2015, 11:e1005058.

67. Khare S, Nagle AS, Biggart A, Lai YH, Liang F, Davis LC, Barnes SW, Mathison CJ, Myburgh E, Gao MY, et al: Proteasome inhibition for treatment of leishmaniasis, Chagas disease and sleeping sickness. Nature 2016, 537:229–233.

68. Wyllie S, Thomas M, Patterson S, Crouch S, De Rycker M, Lowe R, Gresham S, Urbaniak MD, Otto TD, Stojanovski L, et al: Cyclin-dependent kinase 12 is a drug target for visceral leishmaniasis. Nature 2018, 560:192–197.

69. Bjornsti M-A, Knab AM, Benedetti P: YeastSaccharomyces cerevisiae as a model system to study the cytotoxic activity of the antitumor drug camptothecin. Cancer Chemotherapy and Pharmacology 1994, 34:S1–S5.

70. Hoepfner D, McNamara CW, Lim CS, Studer C, Riedl R, Aust T, McCormack SL, Plouffe DM, Meister S, Schuierer S, et al: Selective and specific inhibition of the plasmodium falciparum lysyl-tRNA synthetase by the fungal secondary metabolite cladosporin. Cell Host Microbe 2012, 11:654–663.

71. Lenski RE: Experimental evolution and the dynamics of adaptation and genome evolution in microbial populations. Isme j 2017, 11:2181–2194.

72. Maeda T, Iwasawa J, Kotani H, Sakata N, Kawada M, Horinouchi T, Sakai A, Tanabe K, Furusawa C: High-throughput laboratory evolution reveals evolutionary constraints in Escherichia coli. Nat Commun 2020, 11:5970.

73. Card KJ, Thomas MD, Graves JL, Jr., Barrick JE, Lenski RE: Genomic evolution of antibiotic resistance is contingent on genetic background following a long-term experiment with Escherichia coli. Proc Natl Acad Sci U S A 2021, 118.

74. Rock FL, Mao W, Yaremchuk A, Tukalo M, Crépin T, Zhou H, Zhang YK, Hernandez V, Akama T, Baker SJ, et al: An antifungal agent inhibits an aminoacyl-tRNA synthetase by trapping tRNA in the editing site. Science 2007, 316:1759–1761.

75. Xu Y, Liu F, Chen S, Wu J, Hu Y, Zhu B, Sun Z: In vivo evolution of drug-resistant Mycobacterium tuberculosis in patients during long-term treatment. BMC Genomics 2018, 19:640.

76. Xie SC, Gillett DL, Spillman NJ, Tsu C, Luth MR, Ottilie S, Duffy S, Gould AE, Hales P, Seager BA, et al: Target Validation and Identification of Novel Boronate Inhibitors of the Plasmodium falciparum Proteasome. J Med Chem 2018, 61:10053–10066.

77. Allmeroth K, Horn M, Kroef V, Miethe S, Müller RU, Denzel MS: Bortezomib resistance mutations in PSMB5 determine response to second-generation proteasome inhibitors in multiple myeloma. Leukemia 2021, 35:887–892.

78. Ellis LM, Hicklin DJ: Resistance to Targeted Therapies: Refining Anticancer Therapy in the Era of Molecular Oncology. Clin Cancer Res 2009, 15:7471–7478.

79. Cingolani P, Platts A, Wang le L, Coon M, Nguyen T, Wang L, Land SJ, Lu X, Ruden DM: A program for annotating and predicting the effects of single nucleotide polymorphisms, SnpEff: SNPs in the genome of Drosophila melanogaster strain w1118; iso-2; iso-3. Fly (Austin) 2012, 6:80–92.

80. Robinson JT, Thorvaldsdottir H, Winckler W, Guttman M, Lander ES, Getz G, Mesirov JP: Integrative genomics viewer. Nat Biotechnol 2011, 29:24–26.

81. Benjamini Y, Hochberg Y: Controlling the false discovery rate: a practical and powerful approach to multiple testing. Journal of the Royal statistical society: series B (Methodological) 1995, 57:289–300.

82. Carter H, Douville C, Stenson PD, Cooper DN, Karchin R: Identifying Mendelian disease genes with the variant effect scoring tool. BMC Genomics 2013, 14 Suppl 3:S3.

