## Supplemental Information for "*In vitro* evolution and whole genome analysis to study chemotherapy drug resistance in haploid human cells"

| | | | | |
|---|---|---|---|---|
| $EC_{50}$ curve | 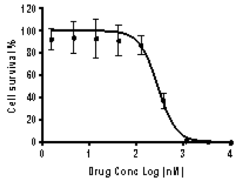 | 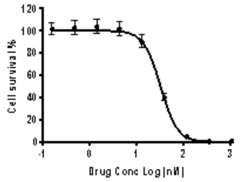 | 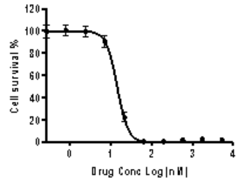 | 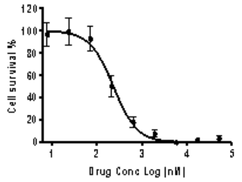 |
| Drug / $EC_{50}$ value | Puromycin 288.44 nM (positive control) | Paclitaxel 19.43 nM | Gemcitabine 34.12 nM | Doxorubicin 95.50 nM |
| $EC_{50}$ curve | 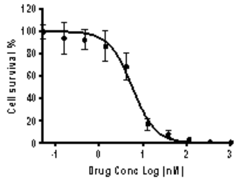 | 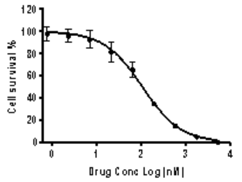 | 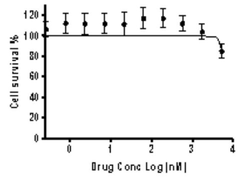 | 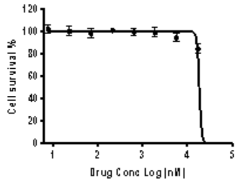 |
| Drug / $EC_{50}$ value | Topotecan 4.81 nM | Etoposide 338.60 nM | Sorafenib > 5 μM | Tamoxifen > 10 μM |
| $EC_{50}$ curve | 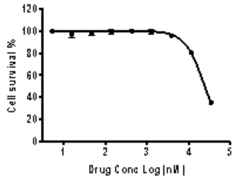 | 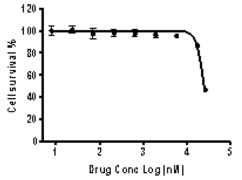 | 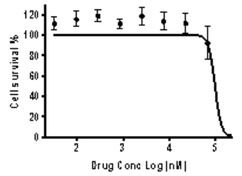 | 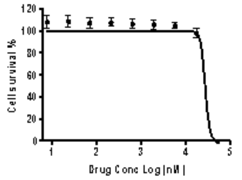 |
| Drug / $EC_{50}$ value | Linsitinib > 10 μM | Everolimus > 10 μM | Imatinib > 10 μM | Rapamycin > 10 μM |
| $EC_{50}$ curve | 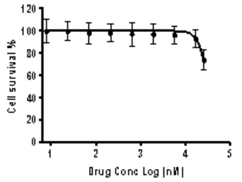 | 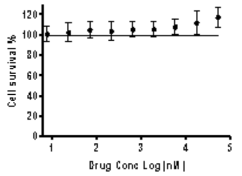 | 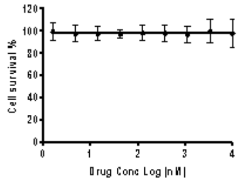 | 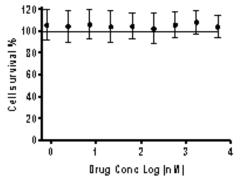 |
| Drug / $EC_{50}$ value | Pterostilbene > 10 μM | Thalidomide N/A | Carboplatin N/A | Imatinib mesylate N/A |

**Table S1. Summary of all drugs tested for IVIEWGA.**

Different anticancer FDA-approved drugs were tested to determine their $EC_{50}$. ATP levels were measured via bioluminescence (CellTiterGlo) to determine drug sensitivity in a dose-response assay for 48h with serial dilutions of the drug (10μM max, 1:3 serial dilutions, 8 technical replicates per concentration point). Those drugs showing $EC_{50}$ values below 1μM were considered for IVIEWGA. Puromycin was used as a positive control.

| | | BR1 | BR2 | BR3 | AVRG | STDEV |
|---|---|---|---|---|---|---|
| DOXORUBICIN | DOX-WT1 | 45.2 | 51.0 | 41.9 | 46.1 | 4.6 |
| | DOX-R1 | 840.9 | 1144 | 945 | 976.6 | 154 |
| | DOX-R2 | 833.7 | 1106 | 957.8 | 965.8 | 136.3 |
| | DOX-R3 | 1132 | 1292 | 1225 | 1216.3 | 80.4 |
| | DOX-WT5 | 150 | 150.6 | 134.2 | 144.9 | 9.3 |
| | DOX-R4a | 1373 | 1403 | 1162 | 1312.7 | 131.3 |
| | DOX-R4b | 503.4 | 504.5 | 466.6 | 491.5 | 21.6 |
| | DOX-R5 | 1510 | 1513 | 1455 | 1492.7 | 32.7 |
| GEMCITABINE | GEM-WT2 | 59.3 | 61.5 | 57.8 | 59.5 | 1.9 |
| | GEM-R1 | 21774 | 22069 | 20377 | 21406.7 | 903.8 |
| | GEM-R2 | 19756 | 19917 | 16721 | 18798 | 1800.5 |
| | GEM-R3 | 27722 | 25236 | 20529 | 24495.7 | 3653.2 |
| | GEM-WT3 | 9.4 | 8.8 | 8.1 | 8.7 | 0.7 |
| | GEM-R4 | 60.1 | 58.9 | 45.3 | 54.8 | 8.2 |
| | GEM-R5 | 82.1 | 78.3 | 66.1 | 75.5 | 8.4 |
| | GEM-R6 | 33.6 | 28.1 | 24.3 | 28.7 | 4.6 |
| PACLITAXEL | PTX-WT4 | 22.8 | 20.6 | 21.2 | 21.5 | 1.2 |
| | PTX-R1 | 120.6 | 231.5 | 251.4 | 201.2 | 70.5 |
| | PTX-R2a | 89.4 | 97.9 | 102.9 | 96.7 | 6.8 |
| | PTX-R2b | 152.9 | 255 | 215.3 | 207.7 | 51.5 |
| | PTX-R3 | 208.9 | 283.9 | 221 | 237.9 | 40.3 |
| | PTX-WT5 | 20.4 | 19.3 | 13 | 17.5 | 4 |
| | PTX-R4 | 855.3 | 959.1 | 843.9 | 886.1 | 63.5 |
| | PTX-R5 | 824.8 | 763.6 | 659.3 | 749.2 | 83.7 |
| | PTX-R6 | 1018 | 1073 | 981.7 | 1024.2 | 46 |
| TOPOTECAN | TPT-WT6 | 5.6 | 5.7 | 5.6 | 5.6 | 0.1 |
| | TPT-R1 | 140.8 | 150.4 | 79.9 | 123.7 | 38.2 |
| | TPT-R2 | 165.7 | 299 | 169.7 | 211.5 | 75.8 |
| | TPT-R3 | 211.2 | 291.2 | 177.2 | 226.5 | 58.5 |
| | TPT-WT7 | 3.4 | 3.7 | 3.6 | 3.6 | 0.2 |
| | TPT-R4a | 34.8 | 23.9 | 29.4 | 29.4 | 7.7 |
| | TPT-R4b | 59.2 | 42.1 | 51.5 | 50.9 | 12.1 |
| | TPT-R4c | 68.9 | 64.9 | 66.9 | 66.9 | 2.8 |
| ETOP. | ETP-WT3 | 292.8 | 363.6 | 359.4 | 338.6 | 39.7 |
| | ETP-R4 | 5876 | 9851 | 8235 | 7987.3 | 1999 |
| | ETP-R5 | 4230 | 5496 | 4010 | 4578.7 | 802 |
| | ETP-R6 | 3742 | 3022 | 2715 | 3159.7 | 527.2 |

**Table S2. $EC_{50}$ values for all drug-specific resistant lines and their isogenic parental cell line (WT).**

$EC_{50}$ averaged values (AVRG) are presented as mean of 4-8 technical replicates with individual biological replicates (BR) overlaid. Doxorubicin (DOX), Gemcitabine (GEM), Paclitaxel (PTX), Topotecan (TPT) and Etoposide (ETP) were used to generate resistant lines as described in methods.

**Table S3. Sequencing sample characteristics and statistics (attached Excel file).**

| Noncoding Variants | |
|---|---|
| *SnpEff Annotation* | *Effect Classification* |
| 3_prime_UTR_variant | Intergenic |
| 5_prime_UTR_premature_start_codon_gain_variant | Intergenic |
| 5_prime_UTR_variant | Intergenic |
| TF binding site variant | Intergenic |
| Conservative inframe deletion | Inframe deletion |
| Conservative inframe insertion | Inframe insertion |
| Disruptive inframe deletion | Disruptive inframe deletion |
| Disruptive inframe insertion | Disruptive inframe insertion |
| Downstream gene variant | Intergenic |
| Intergenic region | Intergenic |
| Intragenic variant | Intragenic |
| Intron variant | Intron |
| Sequence feature | Intergenic |
| Splice acceptor variant & intron variant | Splice region plus intron variant |
| Splice acceptor variant & splice donor variant & splice region variant & 5 prime UTR variant & intron variant | Splice region plus intron variant |
| Splice acceptor variant & splice donor variant & splice region variant & intron variant | Splice region plus intron variant |
| Splice acceptor variant & splice region variant & conservative inframe deletion & intron variant | Splice region plus intron variant |
| Splice acceptor variant & splice region variant & disruptive inframe deletion & intron variant | Splice region plus intron variant |
| Splice donor variant & intron variant | Splice region plus intron variant |
| Splice region variant | Splice region plus intron variant |
| Splice region variant & intron variant | Splice region plus intron variant |
| Splice region variant & synonymous variant | Splice region plus intron variant |
| **Coding Variants** | |
| *SnpEff Annotation* | *Effect Classification* |
| Frameshift variant | Frameshift |
| Frameshift variant & splice acceptor variant & splice donor variant & splice region variant & intron variant | Frameshift |
| Frameshift variant & splice acceptor variant & splice region variant & intron variant | Frameshift |
| Frameshift variant & splice region variant | Frameshift |
| Frameshift variant & stop gained | Frameshift plus stop-gained |
| Missense variant | Missense |
| Missense variant & splice region variant | Missense |
| Protein-protein contact | Other nonsynonymous coding |
| Start lost | Start lost |
| Stop gained | Stop gained |
| Stop lost | Stop lost |
| Structural interaction variant | Other nonsynonymous coding |
| Synonymous variant | Synonymous |

**Table S4. A. Summary of effect classification of the SNVs and Indels from snpEff annotations.**

**Table S5. Summary of mutation types for each individual clone (attached Excel file).**

**Table S6. All SNVs called from the 28 samples (attached Excel file).**

| Gene | Sample | Type | Amino acid change | AF |
|---|---|---|---|---|
| *AC091801.1* | DOX-R4a | MS | His13Asn | 1 |
| *TOP2A* | DOX-R2, DOX-R3 | MS | Pro803Thr | 0.89, 0.87 |
| *EDNRA* | DOX-R2, DOX-R3 | MS | Ala143Ser | 0.86, 0.88 |
| *ITGB3* | DOX-R2, DOX-R3 | MS | Cys627Phe | 0.88, 0.96 |
| *PRAMEF11* | DOX-R2, DOX-R3 | MS | Gln377Lys | 0.91, 0.90 |
| *NCAN* | DOX-R3 | MS | G939Ser | 0.97 |
| *TRIM45* | DOX-R3 | MS | Gln305Lys | 0.89 |
| *SLC13A4* | DOX-R4b | MS | Gly165His | 1 |
| *KCNC3* | DOX-R5 | MS | Ser379Ile | 0.86 |
| *SPG7* | DOX-R5 | MS | Lys593Asn | 1 |
| *WDR87* | DOX-R5 | MS | Arg2418Met | 0.89 |
| *WDR33* | ETP-R3 | MS | Pro622Thr | 1 |
| *STARD9* | GEM-R4, GEM-R5, GEM-R6 | MS | His1021Tyr (R4, R5, R6), Ser1330Ile (R5) | 0.48, 0.47, 0.45, 0.33 |
| *SLCO3A1* | PTX-R2b, PTX-R6 | MS | Ile587Asn (R2b), Ala263Thr (R6) | 0.23, 0.07 |
| *CYP1B1* | TPT-R1, TPT-R4a, TPT-R4b, TPT-R4c | MS | Val432Leu, Asp217Glu (R4a, R4b, R4c) | 0.13, 0.40, 0.43, 0.42 |
| *TOP1* | TPT-R4a, TPT-R4b, TPT-R4c | FS | His81fs | 1 |
| *CEACAM1* | TPT-R4c | MS | Asp479Glu | 1 |
| *VAMP7* | TPT-R4c | MS | Gln57Lys | 1 |
| *ZFP36* | TPT-R4c | MS | Asp260Tyr | 1 |
| *DCK* | GEM-R1, GEM-R2, GEM-R3 | MS, FS | Ser129Tyr (R1, R2), Asn80fs (R1. R3), Asn113fs (R2), Thr184fs (R3) | 0.78, 0.25; 0.28, 0.43; 0.09, 0.14 |
| *GRM3* | ETP-R3 | SIV | - | 1 |

**Table S7. Potential target genes fitting the filtering criteria.**

Listed genes are the completed set of mutations that presented with AF > 0.85, and modified the coding potential, or were mutated multiple times at independent samples. AF, allele frequency; MS, missense; SIV, structural variant; FS, frameshift.

**Table S8. All CNVs called from the 28 samples (attached Excel file).**

| Gene | Matched Type | Drug | Mutated Cell Lines |
|---|---|---|---|
| ABCB1 | CNV | paclitaxel | 70 |
| ABCB4 | CNV | paclitaxel | 66 |
| CYP1B1 | MS | topotecan | 56 |
| DCK | MS | gemcitabine | 11 |
| RAD51 | CNV | etoposide | 1 |
| RRM1 | CNV | gemcitabine | 7 |
| SLC13A4 | MS | doxorubicin | 32 |
| SLCO3A1 | MS | paclitaxel | 32 |
| SPG7 | MS | doxorubicin | 57 |
| TOP1 | CNV | topotecan | 1 |
| TOP1 | FS | topotecan | 4 |
| TOP2A | MS | doxorubicin | 56 |
| WDR33 | MS | etoposide | 79 |
| WNT3A | CNV | etoposide | 42 |
| WNT3A | CNV | topotecan | 42 |
| WWOX | CNV | paclitaxel | 9 |
| WWOX | CNV | topotecan | 9 |

**Table S9. Number of cancer cell lines with matched alterations in implicated resistance genes.**

Mutations in corresponding genes and of a matching type to those observed as drug resistance mutations in HAP-1 were also found in many cancer cell lines with mutation and CNV calls available through the cancer cell dependency map (DepMap) database. CNV, copy-number variant. MS, missense variant. FS, frameshift variant.

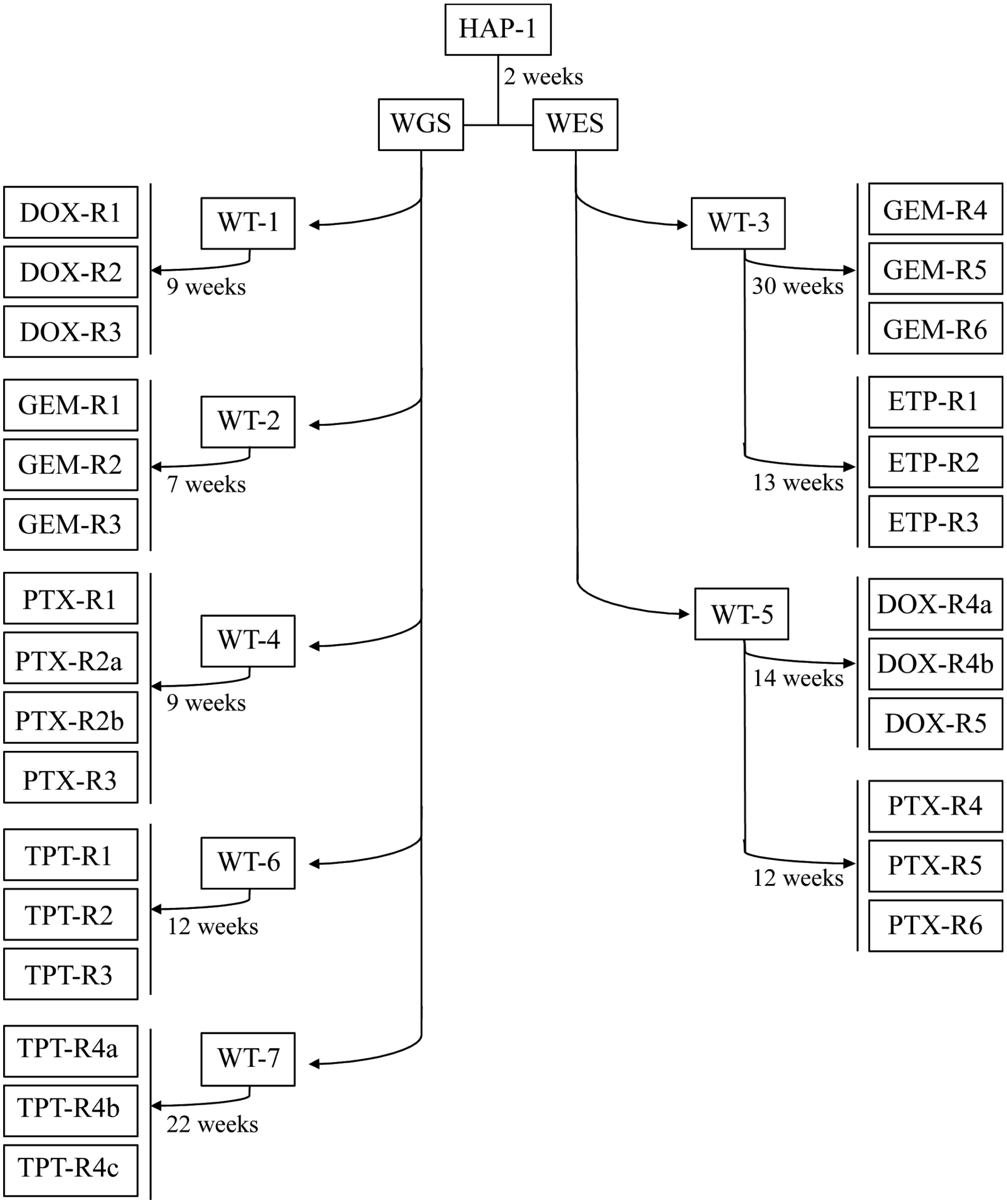

**Figure S1. Hierarchical tree of the parent clones and the different drug-specific replicates.** At least 3 independent HAP-1 drug-resistant clones were generated directly from their isogenic wildtype parents (WT) and their DNA was whole genome (WGS) or whole exome (WES) sequenced. The independent HAP-1 clones were subjected to increasing sublethal concentrations of the drugs based on the starting $EC_{50}$ values for each chemotherapeutic agent until they acquired resistance. The anticancer drugs used for the study were doxorubicin (DOX), gemcitabine (GEM), paclitaxel (PTX), topotecan (TPT) and etoposide (ETP). The time required to generate drug-resistant clones varied depending on the drug from 7 weeks up to 30 weeks (49 to 210 generations). In a few cases, independent selections could not be achieved and dependent clones with a shared lineage were collected.

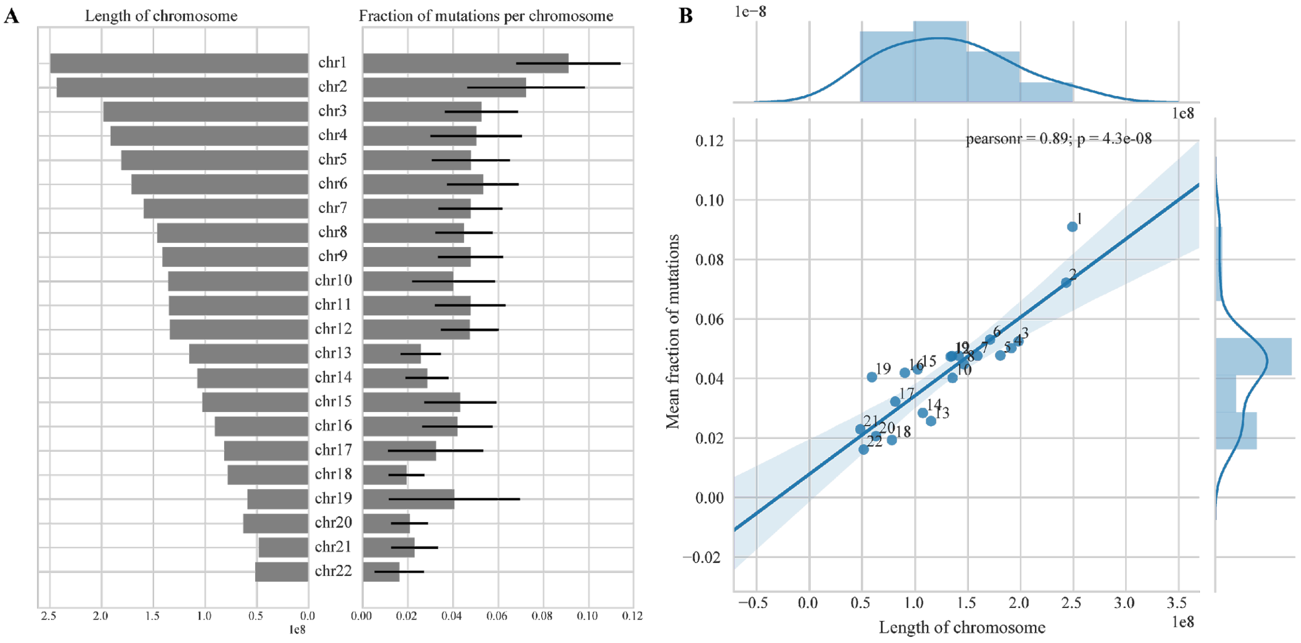

**Figure S2. Near-normal distribution of the mutations in respect to chromosome length.**

A. A bar plot showing length of chromosomes (left) and the fraction of mutations (right) across all samples for each chromosome. Error bars show the standard deviation between samples for their fraction of mutations. B. The length of chromosome (x-axis) shows a 0.89 correlation to the mean fraction of mutations (y-axis) for the samples.

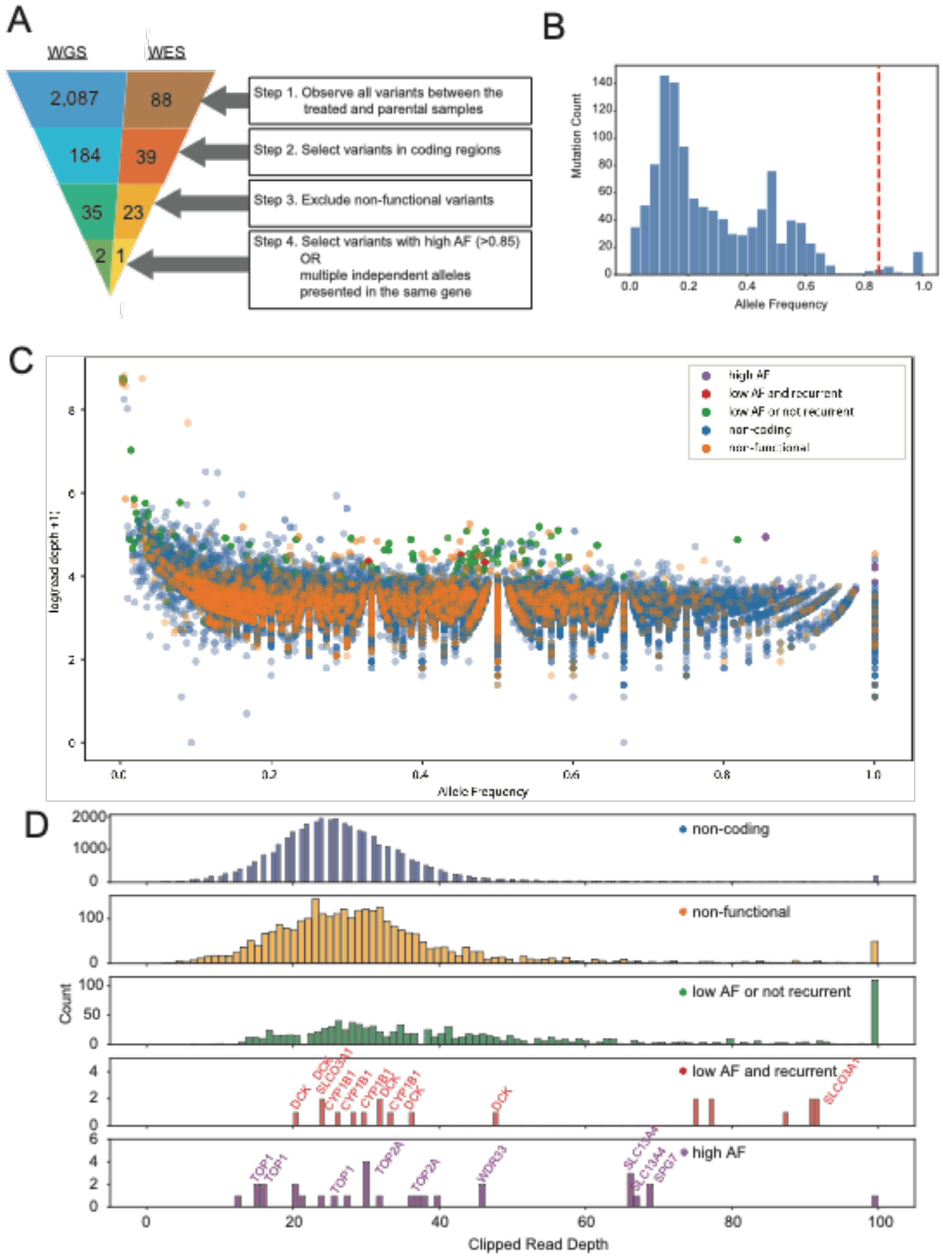

**Figure S3. Mutation filtering pipeline strategies and quantitative summary.**
**A.** Variant filtering pipeline and the average number of mutations per WGS and WES samples. Allelic fraction versus read depth for candidate resistance variants. **B.** Histogram of allelic fractions of coding and functional variants detected in HAP-1 IVIEWGA study. **C.** Scatter plot showing the relationship between read depth and allelic fraction across all variants detected. **D.** Histograms showing the read depth of variants in each category. Variant sites for final selected targets generally had a read depth of 20 or higher.

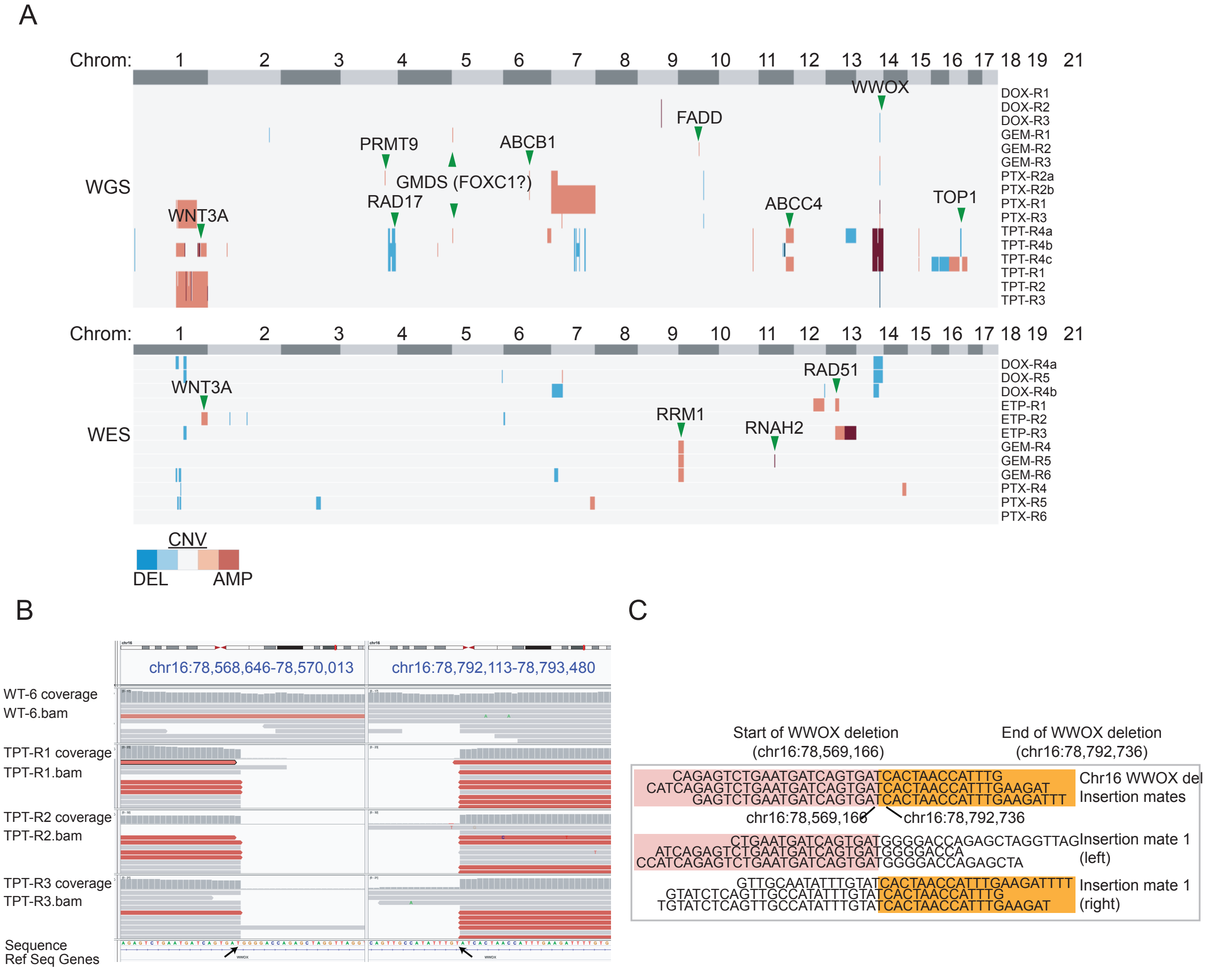

**Figure S4. A. CNV events across samples in each chromosome for WGS and WES samples.** Events less than 0.05% (5E-04) of the chromosome length and events from chromosome X and Y are not shown. The full list of CNV events can be found in Table S8. Candidate genes in small CNVs were identified by examining protein coding genes in minimal amplified intervals to look for classes of genes known to be associated with resistance (ABC transporters, DNA damage response and cell cycle, cytochrome p450s) and performing literature searches on potential candidates. **B.** IGV screen view of the *WWOX* deletion event showing the start (left) and end (right) of the deletion as well as a histogram of coverage at each base. Reads with an abnormal paired-end insert size (~223,000 versus ~200 bases) are highlighted in red. **C.** Portions of reads covering the start and end of the deletion events are shown and allow the identification of the exact breakpoint.

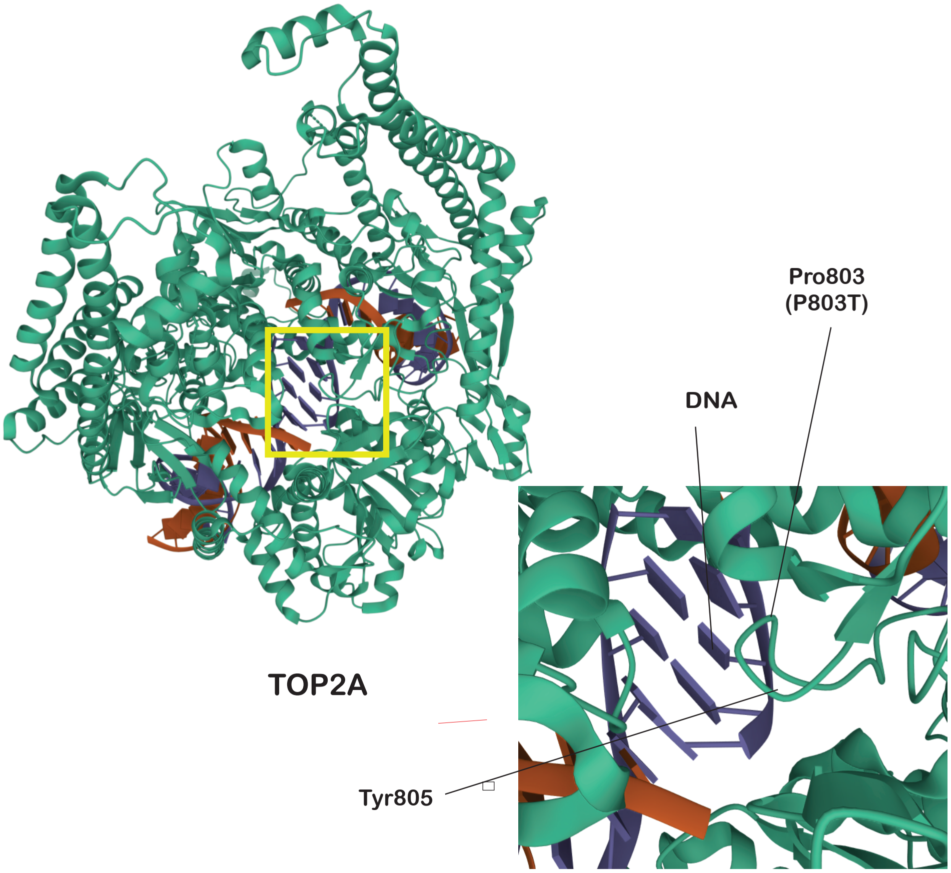

**Figure S5. Crystal structure of *TOP2A*.**
Structures  highlighting (yellow box) the allele Pro803Thr next to the principal drug-binding DNA locus, Tyr805 (structure taken from PDBe 4FM9(Wendorff et al., 2012)).

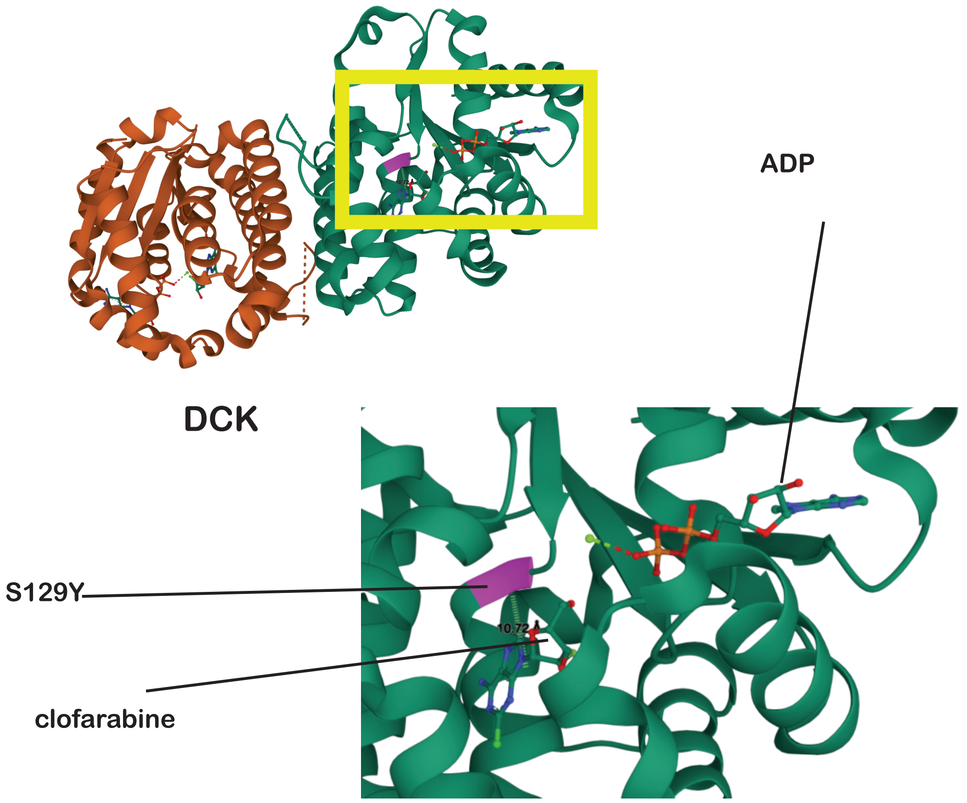

**Figure S6. Crystal structure of deoxycytidine kinase (*DCK*).**
Structure shows the location of the Ser129Tyr (pink) allele near (~10Å) the small molecule inhibitor (clofarabine) and ATP/ADP binding pocket (structure taken from PDBe 2A7Q(Zhang et al., 2006)).

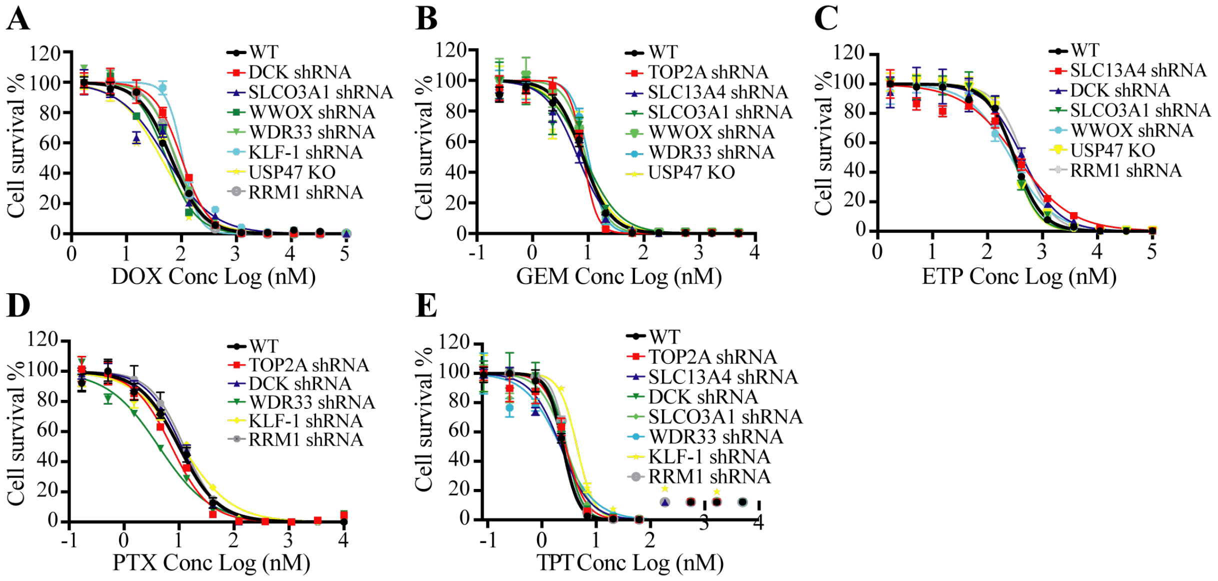

**Figure S7. Cross-drug resistance $EC_{50}$ curves.**

$EC_{50}$ curves for all validated gene candidates that showed absence of multidrug resistance (MDR) pathways against the other drugs used in the study. Target gene candidates tested against: A. DOX, B. GEM, C. ETP, D. PTX, and E. TPT. Experiments were conducted in technical quadruplicates.

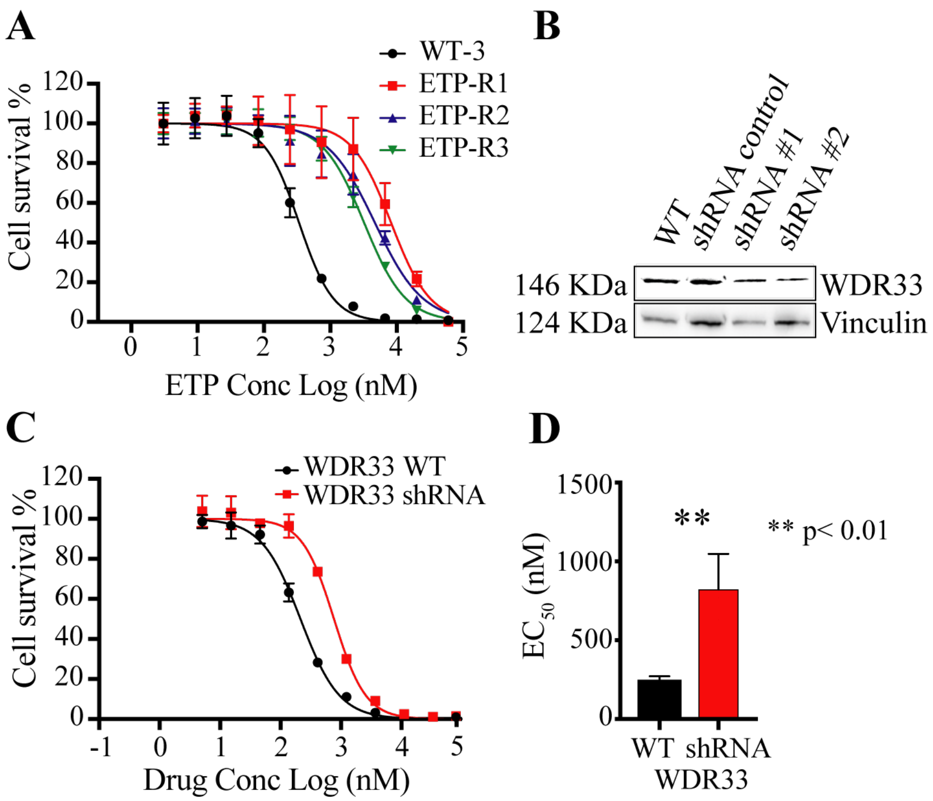

**Figure S8. ETP target genes and validation results.**

A. $EC_{50}$ curves for initial screening ETP resistance. B. Western blot with their gene depletion downregulates protein levels for *WDR33*. C. $EC_{50}$ curves of the WT and shRNA knock-down cell lines. D. Boxplot of the WT and shRNA knockdown cell lines for *WDR33*. Data is represented by mean ± s.e.m. with n=3 biological replicates overlaid and n=4-8 technical replicates. ** = p value < 0.01. p values determined by two-tailed *t* test.

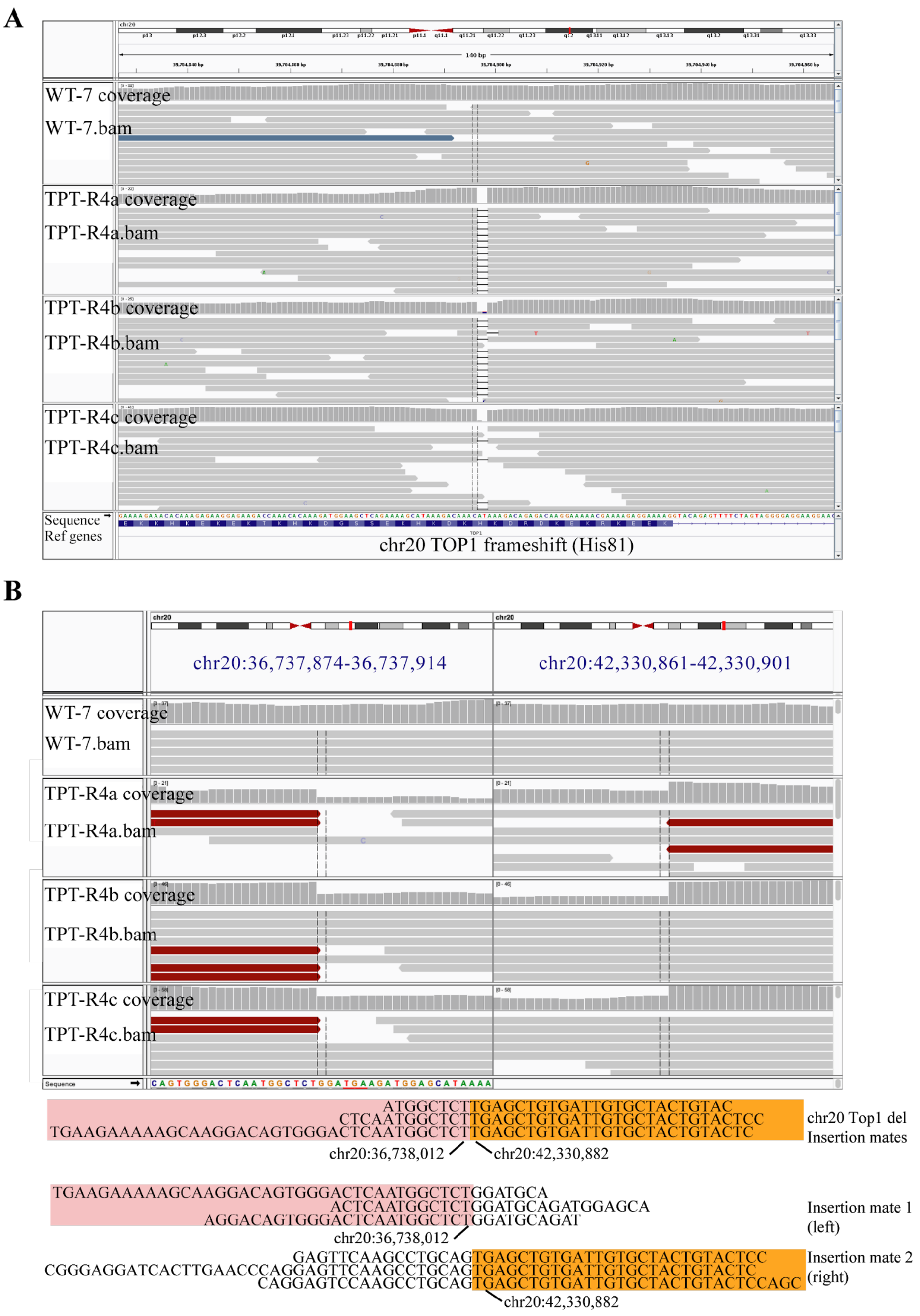

**Figure S9. IGV views for the TPT target genes.**

A. IGV view of *TOP1* frameshift (His81) at TPT cell lines. B. IGV view of the start of the deletion event that includes *TOP1,* where the reads dropped at TPT-R4a, TPT-R4b, and TPT-R4c cell lines compared to WT-7. Left shows the start of the deletion event and right shows the end. At the drop of the reads, there are also insertion events in the replicates but not in the WT.

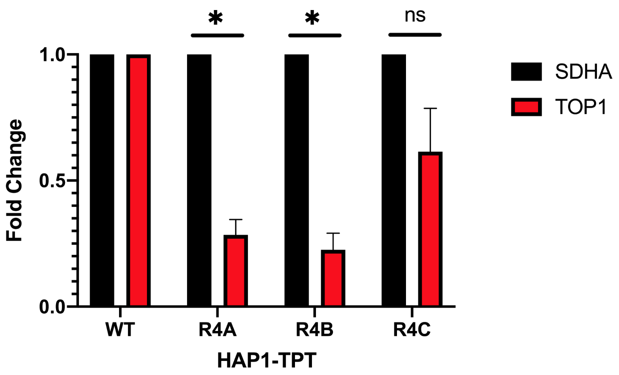

**Figure S10. RT-qPCR quantifying expression of TOP1 in TPT-WT, TPT-R4a-c resistant lines.**

HAP1 wild type and TPT-R4a-c resistant lines were subject to RNA extraction, followed by cDNA generation using random hexamers. Then, qPCR was utilized to measure mRNA expression level of TOP1 in each line relative to the succinate dehydrogenase (SDHA) controls.

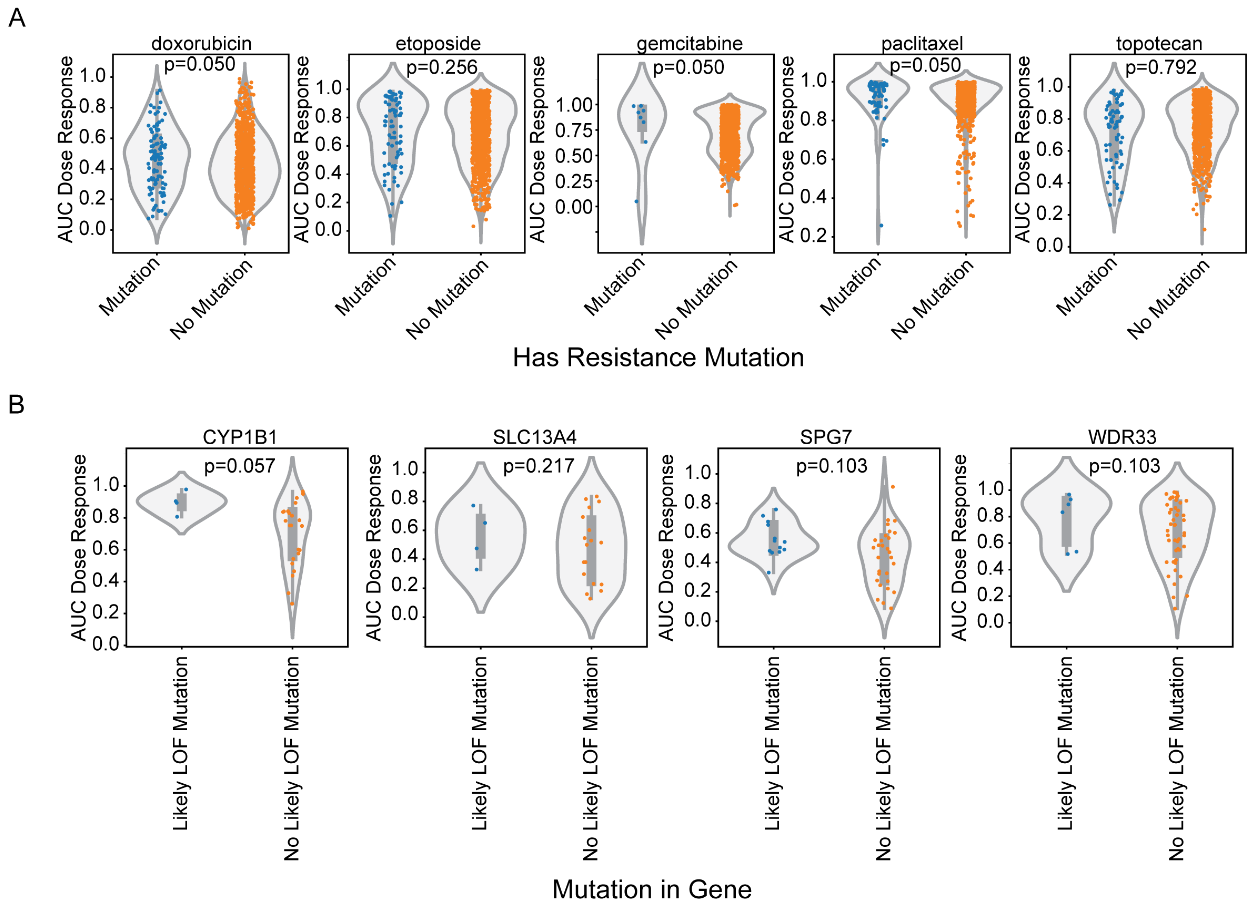

**Figure S11. HAP1 IVIEWGA implicated drug resistance mutations in GDSC cell lines.** **A)** Area under the dose-response curve (AUC) distributions of cell lines from the Genomics of Drug Sensitivity in Cancer (GDSC) dataset treated with the 5 drugs and grouped based on the presence of a matched mutation from Table 2. **B)** AUC distributions of all cell lines carrying a likely loss of function (LOF) SNV based on functional effect predictions by VEST. Silent mutations are included in the "No likely LOF mutation" category. P-values were calculated using a Mann-Whitney U test for each comparison. Benjamini-Hochberg adjusted p-values are shown. SNV, single nucleotide polymorphism.

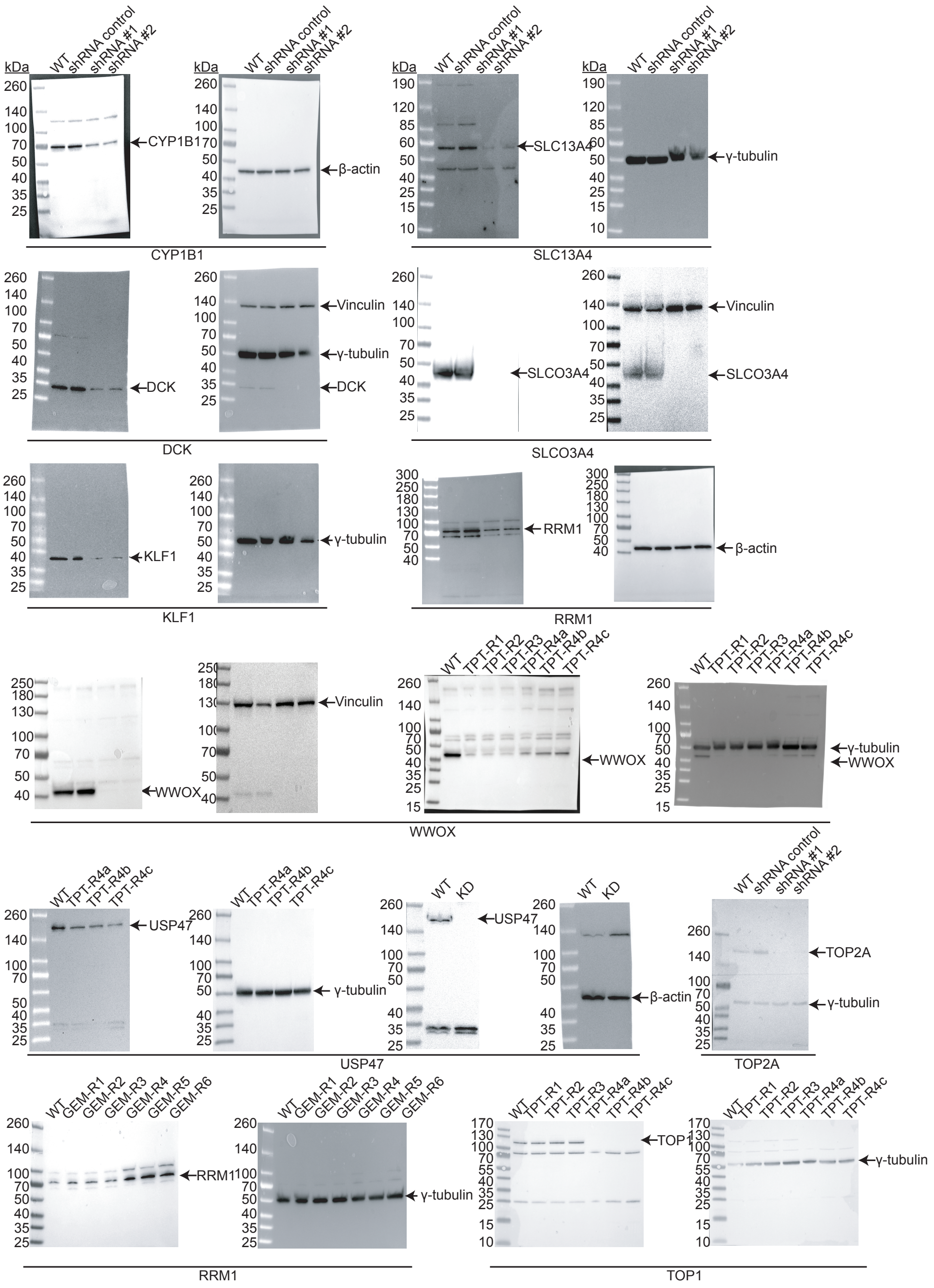

**Figure S12. Western Blots (full membrane) for validated genes.**

Gene depletion via shRNA or CRISPR/Cas9 of all 10 validated genes as well as overexpression or downregulation of the drug-resistant clones compared to their isogenic parents and the scrambled controls are shown in whole membrane together with their corresponding loading controls. Deoxycytidine kinase (*DCK*), Krüppel-Like Factor 1 (*KLF-1*), Topoisomerase II alpha (*TOP2A*), solute carrier transporters (*SLC13A4* and *SLCO3A1*), the ubiquitin-specific peptidase *USP4*7, *WDR33*, tumor suppressor *WWOX*, the catalytic subunit of ribonucleotide reductase *RRM1* and the cytochrome p450 *CYP1B1* were considered in our study. *Vinculin*, *γ-tubulin* and *β-actin* were used as loading controls.
